# The role of repetitive DNA in re-patterning of major rDNA clusters in Lepidoptera

**DOI:** 10.1101/2022.03.26.485928

**Authors:** Martina Dalíková, Irena Provazníková, Jan Provazník, Patrick Grof-Tisza, Adam Pepi, Petr Nguyen

## Abstract

Genes for major ribosomal RNAs (rDNA) are present in multiple copies organized in tandem arrays. Number and position of rDNA loci can change dynamically and their re-patterning is presumably driven by repetitive sequences. We explored a peculiar rDNA organization in several representatives of Lepidoptera with either extremely large or numerous rDNA clusters. We combined molecular cytogenetics with analyses of second and third generation sequencing data to show that rDNA spreads as a transcription unit and reveal association between rDNA and various repeats. Furthermore, we performed comparative long read analyses between the species with derived rDNA distribution and moths with a single rDNA locus, which is considered ancestral. Our results suggest that satellite arrays, rather than mobile elements, facilitate homology-mediated spread of rDNA via either integration of extrachromosomal rDNA circles or ectopic recombination. The latter arguably better explains preferential spread of rDNA into terminal regions of lepidopteran chromosomes as efficiency of ectopic recombination depends on proximity of homologous sequences to telomeres.

## Introduction

Ribosomal RNAs have a central role in ribosome functions in protein synthesis and thus are a cornerstone for life as we know it (Noller et al., 2017). They are shared by all eukaryotes and have been considered the oldest repetitive fraction (Symonová, 2019) as their genes are present in multiple copies organized in tandem arrays. The genes for major ribosomal RNAs (rDNA), i.e. 18S, 5.8S, and 28S, form a transcription unit, in which internal transcribed spacers (ITS 1 and 2) separate individual genes. In eukaryotic genomes (Prokopowich et al., 2003), there are hundreds or even thousands of rDNA units separated by intergenic spacers (IGS) (Long and Dawid, 1980).

Sequences of rRNA genes and their transcribed spacers have been used in taxonomy for species identification (Wu et al., 2015) or to reconstruct phylogenetic relationships (Fiore-Donno et al., 2012). Moreover, thanks to their cluster organization, the rDNA can be easily detected on chromosomes by fluorescent *in situ* hybridization (FISH), which makes it an important marker in cytogenetic studies (Ferretti et al., 2019; Nguyen et al., 2010; Palacios-Gimenez et al., 2013; Provazníková et al., 2021). The rDNA clusters can be present on autosomes, sex chromosomes or even supernumerary chromosomes, i.e. B chromosomes (Cabral-de-Mello et al., 2011; Poletto et al., 2010; Provazníková et al., 2021; Silva et al., 2014; Zrzavá et al., 2018). While most animal species have only one rDNA locus, up to tens of loci were reported in some extreme cases (Eickbush and Eickbush, 2007; Sochorová et al., 2018 and references therein). The active loci are also called the nucleolar organizer regions (NORs) (Ingle et al., 1975; Kobayashi, 2011), as transcription of major rDNA genes and processing of primary transcripts give rise to a sub-nuclear compartment known as a nucleolus (reviewed in Eickbush and Eickbush, 2007). In general, changes in distribution of rDNA genes are dynamic and rDNA was thus compared to mobile elements (MEs), which, in turn, have been considered an important driver in rDNA re-patterning (Cabral-de-Mello et al., 2011; de Sene et al., 2015; Elliott et al., 2013; Ferretti et al., 2019; Scacchetti et al., 2012).

The order Lepidoptera with its 160,000 species of moths and butterflies represents one of the largest insect radiations (Van Nieukerken et al., 2011). Their rich species and ecological diversity contrast with their conserved genome architecture with the ancestral and the most common chromosome number being n=31 (Ahola et al., 2014; Robinson, 1971; Van’t Hof et al., 2013). Detailed analyses of advanced ditrysian species, such as the peppered moth (*Biston* betularia, n=31; Van’t Hof et al., 2013), the Glanville fritillary (*Melitaea cinxia*, n=31; Ahola et al., 2014), and the tobacco cutworm (*Spodoptera litura*, n=31; Cheng et al., 2017) showed highly conserved synteny and order of genes between homoeologous chromosomes. A typical lepidopteran mitotic complement consists of small dot-shape chromosomes (Fuková et al., 2005; Mediouni et al., 2004; Prins and Saitoh, 2003), which lack localized centromere, i.e. they are holokinetic (Wolf et al., 1997). Moreover, traditional bending techniques failed to differentiate individual chromosomes, which has made the classic cytogenetic research in Lepidoptera rather challenging (Bedo, 1984) and limited it, for long time, only to chromosome counting (Lukhtanov, 2015; Robinson, 1971). However, the use of various FISH modifications provided great insight into evolution of lepidopteran karyotypes (Van’t Hof et al., 2013; Yasukochi et al., 2011), sex chromosomes (Carabajal Paladino et al., 2019; Martina Dalíková et al., 2017a; Šíchová et al., 2015; Vítková et al., 2007), repetitive sequences (Šíchová et al., 2015, 2013), and gene families such as major rDNA (Nguyen et al., 2010; Provazníková et al., 2021).

Number and localization of rDNA loci were determined using FISH with the 18S rRNA probe in various species sampled across Lepidoptera (Fuková et al., 2005; Nguyen et al., 2010; Provazníková et al., 2021; Šíchová et al., 2016, 2015, 2013; Vershinina et al., 2015; Zrzavá et al., 2018). The results implied that one terminal rDNA cluster is probably the ancestral state as it was found across all Lepidoptera. In some ditrysian families, such as Noctuidae and Erebidae, the rDNA cluster moved to interstitial position, which was conserved in all studied species. When multiple clusters are present, they are located terminally in majority of species. Higher numbers of rDNA clusters were observed in representatives of the families Psychidae (3-4 clusters) and Nymphalidae (7-11 clusters), *Biston betularia* (3 clusters, Geometridae) and *Hyalophora cecropia* (3 clusters, Saturniidae), all having the ancestral haploid chromosome number n=31. Thus, spread of rDNA clusters is not clearly associated with large scale chromosome rearrangements such as chromosome fissions or fusions. Unusual distribution of rDNA was observed in the ghost moth, *Hepialus humuli* (Hepialidae), and the horse chestnut leaf miner, *Cameraria ohridella* (Gracillariidae), in which signal of the 18S rDNA probe covered about one half and one fourth of a single NOR-bearing chromosome, respectively (Provazníková et al., 2021). It was proposed that the dynamic rDNA repatterning is due to ectopic recombination, i.e. recombination between non-homologous regions mediated by ubiquitous repetitive sequences (Nguyen et al., 2010). However, the hypothesis is yet to be tested.

In this study, we decided to explore a peculiar rDNA organization in *H. humuli* and *C. ohridella*, with extremely large rDNA clusters and nymphalids *Aglais urticae* and *Inachis io* with seven and eleven loci per haploid genome, respectively. We performed FISH with probes for 18S and 28S rDNA to test whether genes for major rRNAs spread individually or as a transcription unit. Further, we sequenced genomes of all four species and analysed repetitive sequences and their co-localization with rDNA using the RepeatExplorer pipeline (Novák et al., 2013, 2010). We estimated portion of rDNA units associated with identified repeats by analyses of coverage. The co-localization of several repetitive sequences with rDNA was verified by FISH and in *H. humuli* and the nymphalids also by analysis of long reads. The long reads analysis was further performed also in *Phymatopus californicus* (Hepialidae), to compare it with the peculiar *H. humuli* rDNA organization, and in *Lymantria dispar* (Erebidae), *Spodoptera frugiperda* (Noctuidae), and *Plutella xylostella* (Plutellidae) to compare rDNA composition between species with an ancestral and highly derived rDNA distribution. Our work shows that combining molecular cytogenetic techniques with next generation sequencing technologies represent a powerful tool to study evolution of genome architecture in Lepidoptera.

## Results

### FISH with 18S and 28S rDNA probes

To examine the organization of the rDNA clusters in genomes of four studied species, namely: *H. humuli*, *C. ohridella*, *A. urticae*, *and I. io*, FISH with 18S and 28S rDNA gene probes was carried out. Hybridization patterns of 18S rDNA probe of all four species correspond to previous results (Nguyen et al., 2010; Provazníková et al., 2021). Moreover, 28S rDNA probe colocalized with 18S rDNA probe in all cases which suggests that the observed patterns of rDNA distribution are due to spread of the whole rDNA unit. One large major rDNA cluster covering large portion of one chromosomal bivalent was observed in pachytene nuclei of *H. humuli* (Figure 1a) and *C. ohridella* (Figure 1b). Additionally, a strong DAPI-positive heterochromatin block colocalized with major rDNA cluster in *H. humuli* (Figure 1a detail). In pachytene nuclei of *A. urticae* and *I. io*, multiple rDNA clusters were observed as expected. Seven small terminal clusters in pachytene nucleus of *A. urticae* did not colocalize with any heterochromatin blocks (Figure 1c), whereas in pachytene nucleus of *I. io*, 11 terminal signals of various sizes were detected and 6 of them colocalizing with small DAPI-positive blocks (example in Figure 1d detail). Numerous chromosomal bivalents bearing small terminal heterochromatin blocks seem to be typical feature for *I. io* karyotype.

**Figure 1:**
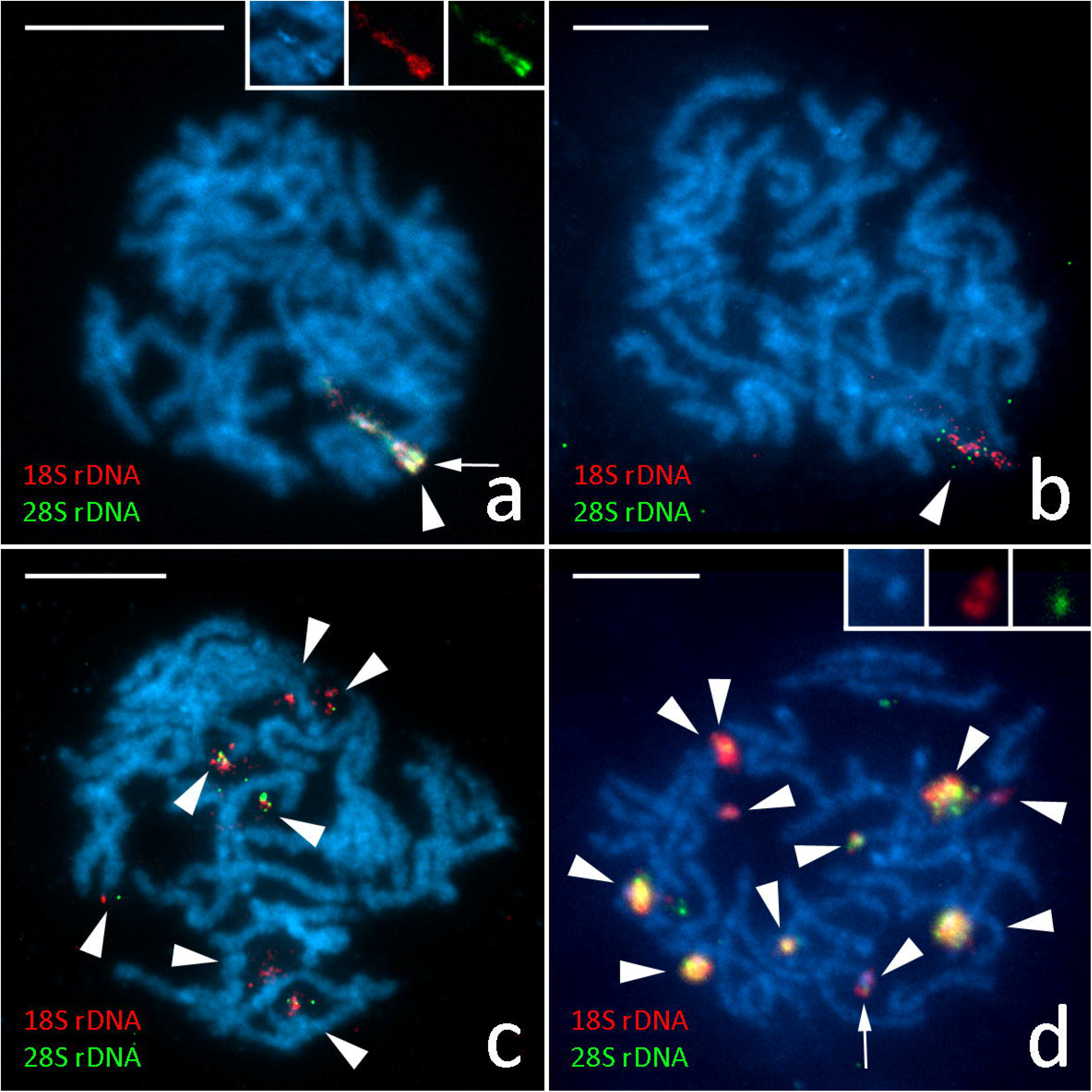
Major rDNA clusters detected by fluorescence in situ hybridization (FISH) on pachytene nuclei of studied species. The 18S rDNA probe (red) and 28S rDNA probe (green), chromosomes are counterstained with DAPI (blue). Major rDNA clusters indicated by arrowheads. **a** – male pachytene nucleus of *H. humuli* (Hepialidae) with detail of the major rDNA cluster colocalizing with heterochromatin blocks in the inset, **b** – male pachytene nucleus of *C. ohridella* (Gracillariidae), **c** – male pachytene nucleus of *A. urticae* (Nymphalidae), **d** – female pachytene nucleus of *I. io* (Nymphalidae) with detail of one of major rDNA clusters colocalizing with a small heterochromatin block in the inset. * clusters colocalizing with DAPI positive heterochromatin. Scale 10 μm

### Repeat Explorer analysis

To identify repeats associated with 45S rDNA we performed Repeat Explorer (RE) analysis in all four studied species. Repetitive sequences with frequent colocalization in the genome can be identified through RE analysis as clusters connected by pair-end reads forming the so-called superclusters. In *C. ohridella*, major rDNA genes were split into two clusters. Surprisingly, these clusters have no connection neither between each other nor to any other identified repeat. The estimation of genome proportion formed by major rDNA in this species is about 0.04 % (Suppl. Table S1). In *I. io*, clusters annotated as 45S rDNA were part of supercluster 11. This supercluster was formed by 3 clusters which were annotated as 28S and LINE R2 element (cluster 27), 18S and 5.8S (cluster 35), and putative satellite (IiSat, cluster 37) with predicted monomer length 157 bp (Suppl. Table S1). As cluster 27 was formed by both 28S rRNA gene and LINE R2 elements (IiR2), genome proportion of major rDNA cannot be determined with certainty from the RE results alone; but these genes could comprise 0.12-0.29% of *I. io* genome (Suppl. Table S1). In *A. urticae*, genes for major rDNA were also divided into several clusters, three clusters 17, 21 and 29 annotated as 45S rDNA formed one supercluster 8 and were not connected to any other repeat by 10 or more shared pair-end reads (Suppl. Table S1). However, after further inspection one contig corresponding to cluster 29 contained tandemly repeated sequence suggesting that a satellite repeat (AuSat) with monomer approx. 400 bp is part of this cluster. Based on RE estimate, the major rDNA clusters formed 0.59 % of *A. urticae* genome. In *H. humuli*, major rDNA genes represented 0.17% of the genome and were all comprised in the cluster 53 which was part of the supercluster 24 together with the cluster 67 annotated as ME from Ty3/Gypsy group (Hh Ty3/gypsyA; Suppl. Table S1).

### FISH with 18S rDNA and ME probes

To verify the results obtained from RE analysis, we mapped ME probe together with 18S probe on chromosome preparations of *H. humuli* and *I. io* by double TSA FISH. In *H. humuli*, both the 18S rDNA probe and the pooled Hh Ty3/GypsyA *RT*, *INT*, and *RH* probes hybridized to the major rDNA cluster region (Figure 2a) and thus confirmed association between Hh Ty3/GypsyA and the major rDNA. Similar pattern was observed in *I. io*, in which both the 18S rDNA probe and the probe for IiR2 *RT* hybridized to all 11 clusters of major rDNA in pachytene nuclei (Figure 2b), thus proving an association between rDNA and the IiR2 ME. Hybridization of the IiSat probe did not provide any clear signal which was probably caused by low quality of generated probe. Additionally, to investigate whether IiR2 is also present in genome of closely related *A. urticae*, we hybridized the 18S rDNA probe and the IiR2 probe to pachytene nuclei of *A. urticae*. However, no clear hybridization signal was observed (results not shown).

**Figure 2:**
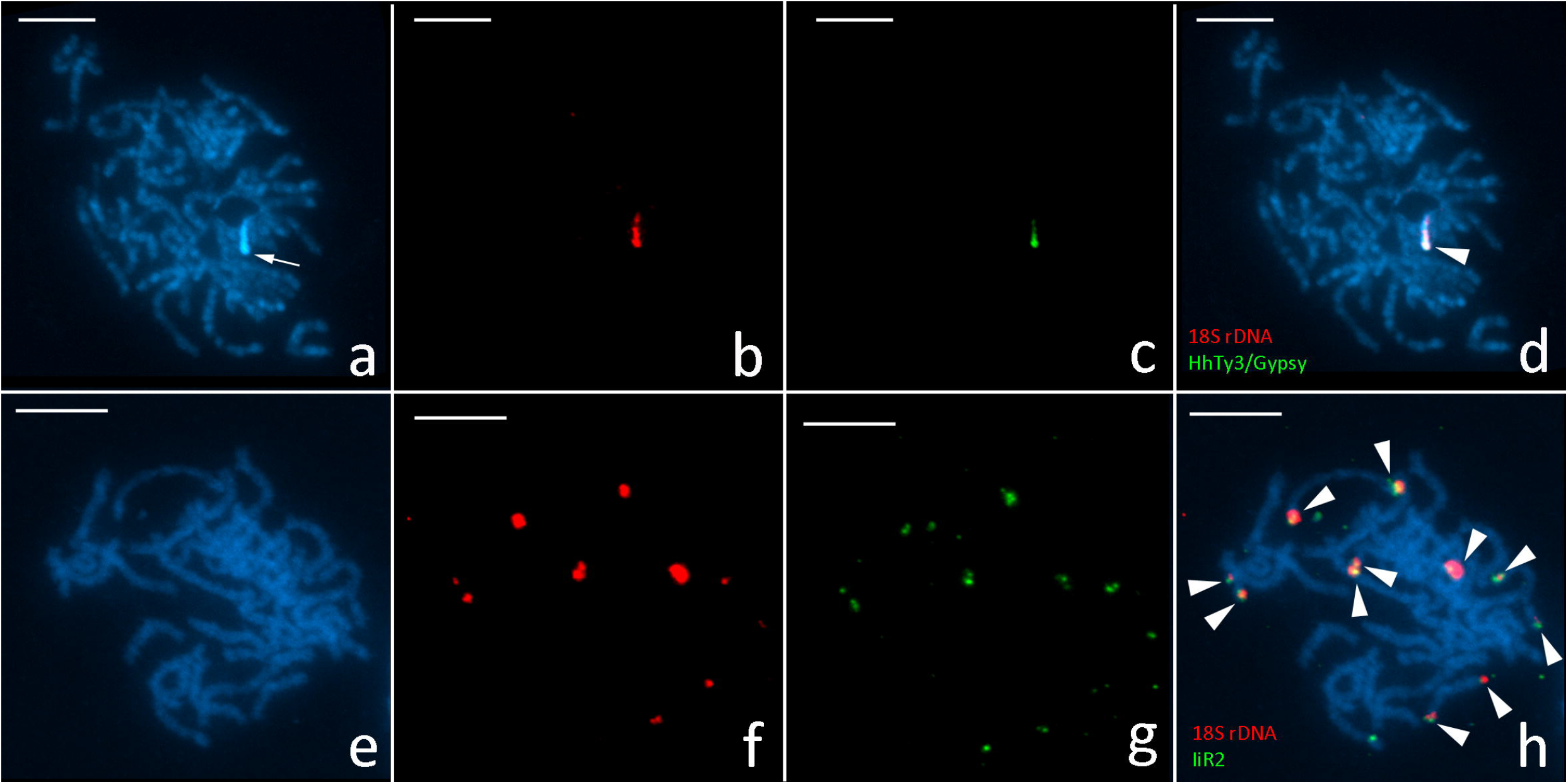
Co-localization of major rDNA and ME sequences of interest as detected by fluorescence *in situ* hybridization (FISH) on pachytene nuclei of studied species. 18S rDNA probe in red (**b, d, f, h**) and HhTy3/GypsyA (**c, d**) and IiR2 probes (**g, h**) in green, chromosomes are counterstained with DAPI (blue; **a**, **d, e, h**). Hybridization signals indicated by arrowhead. **a-d** – male pachytene nucleus of *H. humuli* (Hepialidae), **e-h** – female pachytene nucleus of *I. io* (Nymphalidae). Arrow indicates DAPI positive heterochromatin. Scale 10 μm

### Long read analysis

Long read analysis was used to further verify connection between rDNA and repeats revealed by RE and FISH results. Output of the *H. humuli* sequencing run was poor both in overall yield and read length. After default quality filtering, which was part of the base calling process we obtained 4 Gb in reads with N50 length 6 kb. Due to low coverage of obtained Nanopore data we were able to analyse only 567 reads longer than 15 kb with mean quality (Q) > 10 containing major rRNA genes in *H. humuli*. Most of these reads contained non-functional short copies of three Ty3/Gypsy elements, two LINE elements (from L2 and RTE-RTE groups), two PIF elements (Harbinger and Spy group), and a P element in the IGSs. The organisation and length of the IGSs were highly conserved. Only 18 reads bearing rDNA did not contain any of the mentioned MEs and around 12 % of the reads exhibited some irregularities in the observed pattern (Suppl. Figure S1). All reads containing rDNA were used to assemble rDNA unit including IGS. The resulting Flye assembly contained single circular contig 77 920 bp long corresponding to two complete rDNA units with their lengths differing only in 6bp. Although only one of the observed Ty3/Gypsy elements, Hh Ty3/gypsyA, was detected by the RE analysis as a part of the supercluster containing rDNA, upon careful examination of RE results, all the IGS repeats were connected by shared pair-end reads. Yet these did not suffice to bind rDNA and all associated MEs in one supercluster (Suppl. Table S1).

To test for the presence of such complex IGS in other representative of the family Hepialidae, we have analysed also Nanopore reads of *P. californicus*. After basecalling with default quality filtration, we obtained 40 Gb of data with read length N50 of 22 kb. Total of 596 reads bearing major rDNA passed the filtering for length > 15 kb and Q > 10. Yet in this case, the IGSs were much smaller and contained only ca. 800 bp long microsatellite region (Suppl. Figure S2) consisting of sequence complementary to insect telomeric repeat TTAGG and 231 bp long region of TTATG microsatellite. The Flye assembly yielded a single 30 616 bp long circular contig. However, this contig corresponded to three complete major rDNA units which differed in length of microsatellite region by up to four TTAGG repeats. The major rDNA unit in *P. californicus* is thus about 10 kb long including IGS region.

Due to high coverage of available HiFi PacBio reads of *I. io*, we were able to analyse 3 276 reads containing major rDNA longer than 15 kbp (Suppl. Figure S3). Of these, 2 625 contained at least 200 bp of the satellite recovered by the RE analysis in their IGS. However, the individual major rDNA units differ in length of this satellite array (Suppl. Table S2). In 946 reads, the R2 element was inserted in major rDNA genes. Surprisingly some of these insertions were not limited to the 28S rRNA gene, suggesting ongoing degeneration of rDNA units via R2 insertions in *I. io*. The attempt to assemble the most prevalent variant of complete rDNA unit in *I. io* failed as all assemblies contained more than 300 contigs of variable length, both linear and circular. However, most of the contigs had very low coverage. Moreover, only 27 contigs contained more than 200 bp of any rRNA gene. Out of all obtained contigs, only three had mean coverage over 10% of used PacBio reads and their length varied from aprox. 15.8-33 kb. These three contigs all contained at least some of the major rRNA genes either with or without R2 element insertion and the satellite array with variable length from 4.5 kb to 17 kb. This further emphasizes the variability in IGSs in this species.

In *A. urticae*, the available HiFi PacBio data contained 2 921 reads bearing major rDNA > 15 kb. The long read analysis confirmed overall lack of MEs associated with rDNA as (Suppl. Figure S4) only 179 reads contained either LINE or Ty3/Gypsy element adjacent to major rRNA genes, including R2 element inserted into 28S rDNA. Despite the absence of MEs in IGS of *A. urticae*, this region seems to vary in length between major rDNA units (Suppl. Figure S4). The variation of IGS was reflected also in assembly results as any assembly produced by Flye contained over 70 linear contigs in the length ranging 19 – 69 kb. However, only six contigs contained more than 200 bp of major rDNA and only one of these contigs had mean coverage over 10% of input reads. This contig is 31 kbp long and contains two complete rDNA units including IGS regions. One unit contains R2 insertion in 28S gene and based on dot plot both IGSs contain approx. 2kb of satellite (AuSat) array. Both AuSat IGS satellite arrays contain 6 monomers with variable length from 252 to 408 bp with the last monomer being the shortest one. This satellite was present in most major rDNA units as it was found in 2805 reads containing major rDNA (Suppl. Figure S4). After further inspection of previous RE results in this species, a sequence homologous to the satellite was found among contigs belonging to the cluster 29 (Suppl. Table S1) which also contained a part of the 28S rRNA gene. Interestingly, our RE results did not contain any cluster with sequence homologues to the AuR2 element found in long reads. Considering that samples for RE were sampled from the Czech *A. urticae* population while specimen sequenced by PacBio originated from Great Britain, the R2 insertion into rDNA units may represent inter-population variation in this species.

To test if the variable and/or long IGSs are connected with atypical rDNA genomic organization we performed long read analysis also in species with one major rDNA locus per haploid genome which is supposedly ancestral in Lepidoptera (Nguyen et al., 2010; Provazníková et al., 2021), namely in *Plutella xylostella*, *Spodoptera frugiperda* and *Limatria dispar*. In *P. xylostella*, we analysed 419 PacBio reads containing major rDNA with length at least 15 kb. The rDNA was not associated with any ME, however the IGS contained 850 bp satellite (PxSat) region (Suppl. Figure S5). This array consisted of four monomers with slightly variable length between 248-258 bp with the last monomer being incomplete and only 91 bp long. At least 200 bp of this satellite array was found in 343 reads out of the analysed reads containing major rDNA. The Flye assembly of all filtered rDNA bearing PacBio reads yielded one circular contig approximately 21 kb long. This contig consists of two similarly long complete rDNA units (10.7 and 10.9 kb) which differ in the PxSat array length by 242 bp.

In *L. dispar*, 242 quality filtered PacBio reads contained major rDNA. Surprisingly, major rDNA in this species was associated with two different MEs from LINE R1 group specific for rDNA (Suppl. Figure S6). 51 analysed reads contained at least one of those elements and 8 reads contained both transposons. Flye assembly contained 2 linear contigs 11 kbp and 9.4 kb long with the latter having approx. 10x higher coverage. Neither of these contigs contained complete major rDNA unit, the longer contig contained both R1 elements and the shorter one incomplete major rDNA unit with IGS but only partial 28S rRNA gene.

In *S. frugiperda*, we analysed 115 quality and length filtered PacBio reads containing major rDNA. There were no MEs or satellite sequences observed in these reads (Suppl. Figure S7). Flye assembly consisted of only one circular contig 17.9 kb long which contained two identical complete major rDNA units.

Satellite DNA arrays contained in IGSs of *P. xylostella* representing the ancestral rDNA distribution and both nymphalid species with multiple rDNA clusters seemed to vary in length. Thus, we further characterised these satellite arrays. All three species have similar most represented array length in the PacBio reads, as the medians are ranging from 1.74 to 2.21 kb (Suppl. Table S2, Suppl. Figure S8). However, they differ greatly in maximal observed length, which was 4.67 kb in *P. xylostella* but over 15kb in both nymphalid species (Suppl. Table S2, Suppl. Figure S8). Similar differences can be seen in the satellite array length in the rDNA assemblies. In nymphalid species we obtained multiple contigs with variable satellite length ranging from less than 1 kb to 2.23 kb in *A. urticae* and over 17 kb in *I. io*. (Suppl. Table S2). Whereas, in *P. xylostella* we obtained just one contig with two satellite arrays differing in length by just 242 bp (see above). These results suggest higher variation in length of IGS satellite arrays in *A. urticae* and *I. io* compared to *P. xylostella* (Suppl. Table S2, Suppl. Figure S8). Moreover, the three species differed in the presence of PacBio reads containing satellite sequence without any part the rDNA unit. While we found no such reads in *P. xylostella*, 11 and 30 reads were found in *A. urticae* and *I. io* PacBio data, respectively. As the lengths of PacBio reads bearing only satellite in both nymphalid species exceed the lengths of observed satellite arrays in rDNA assemblies (Suppl. Table S2), these reads either come from very large IGS satellite arrays or they may represent satellite arrays outside the rDNA cluster. The latter is supported by the recently published *A. urticae* genome (Bishop et al., 2021), which contains only 6 terminal rDNA clusters, however, AuSat is found both within the IGS region and right outside two NORs (Suppl. Figure S9).

### Coverage analysis

Paired-end reads produced by Illumina sequencing provided us with sufficient coverage to compare per base abundance of reads aligned to the consensus sequences of the whole rDNA unit of *H. humuli* obtained via long reads analysis (see above) and the most represented complete rDNA unit of *I. io* and *A. urticae* from rDNA assembled contigs (see above). In case of *H. humuli* (Figure 3a, Suppl. Table S3), the repetitive elements associated with major rDNA are most likely present elsewhere in the genome as we observed a uniform coverage of the rDNA genes and varying but higher coverage of the intergenic MEs. In *I. io*, we observed that the R2 element is only present in roughly one third of the copies of the 28S rRNA gene (Figure 3b, Suppl. Tab. S3). The coverage of IiSat region was approx. two times larger compared to rRNA genes (Figure 3b, Suppl. Tab. S3), which similarly to the results obtained from PacBio reads suggest the IiSat presence outside of rDNA clusters and/or the variable length of this satellite array inside IGS regions. Surprisingly, in *A. urticae* all the rDNA unit elements showed even coverage including the AuSat region (Figure 3c, Suppl. Tab. S3). This discrepancy between results obtained through illumina and PacBio reads may represent another inter-populational variability in this species in the repeat content (see above).

**Figure 3:**
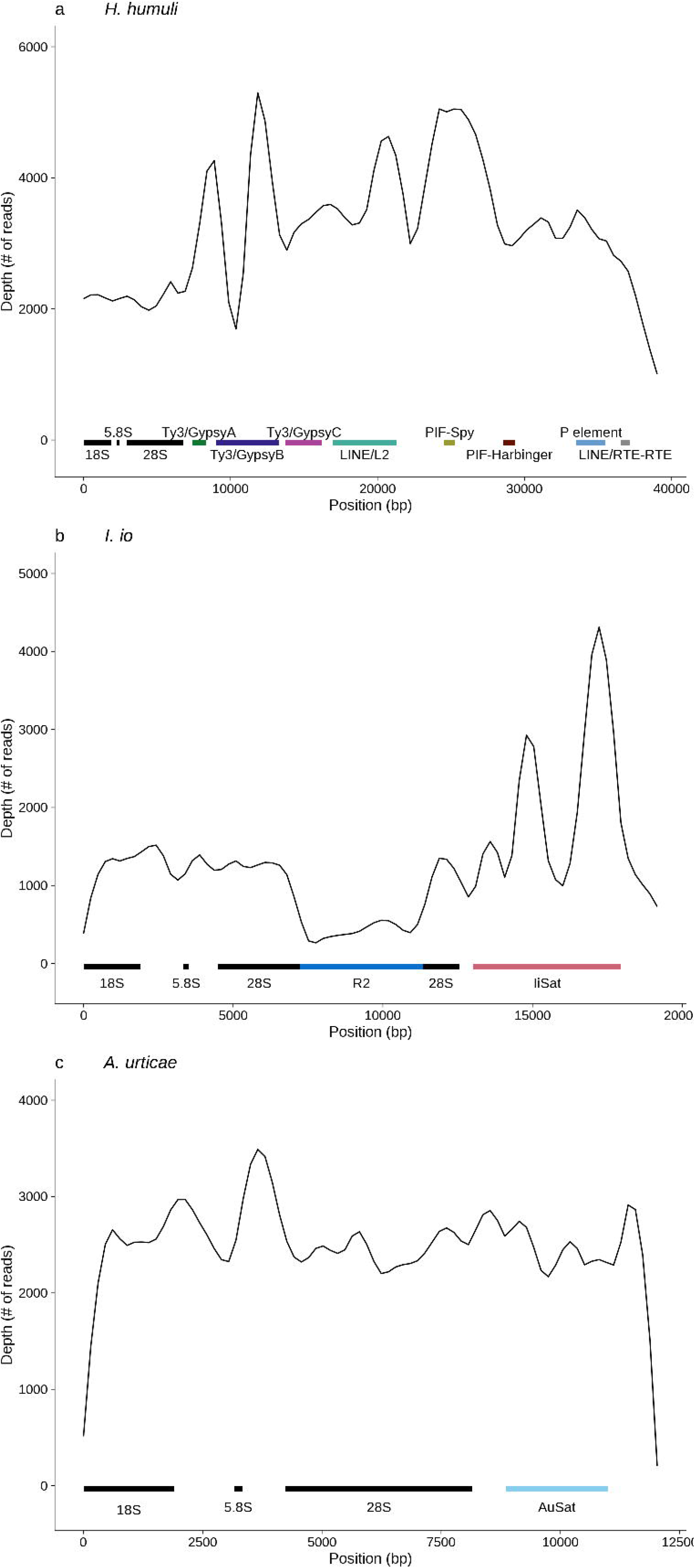
Coverage plot of rDNA units in *H. humuli* (**a**), *I. io* (**b**), and *A. urticae* (**c**). Smoothed (LOESS) counts of aligned sequencing reads for each nucleotide position of the major rDNA cluster. Coloured bars on the bottom represent regions of repetitive elements and position of rDNA genes.

## Discussion

The arrays of major rRNA genes have become a very popular cytogenetic marker in comparative studies of karyotype evolution. Distribution of major rDNA can be relatively stable or rather dynamic in various taxa (Cabral-de-Mello et al., 2011; García-Souto et al., 2016; Nguyen et al., 2010; Perumal et al., 2017) and even intraspecific variability was observed (Baumgärtner et al., 2014; Ferretti et al., 2019; Šíchová et al., 2015). Changes in number and localization of rDNA loci have been ascribed to sequence homogeneity maintained by gene conversion (reviewed in Eickbush and Eickbush, 2007) and chromosomal rearrangements mediated by ectopic recombination (Cabral-de-Mello et al., 2011; Ferretti et al., 2019; Nguyen et al., 2010), transposition (Raskina et al., 2008, 2004) or integration of extrachromosomal circular rDNA (ecc-rDNA; Proux-Wéra et al., 2013).

To gain some insight into mechanism of spread of major rDNA in genomes of moths and butterflies, we examined four lepidopteran species with peculiar distributions of 18S rDNA (Nguyen et al., 2010; Provazníková et al., 2021) namely *H. humuli* and *C. ohridella* with large rDNA locus covering up to half of the chromosome, and *A. urticae* and *I. io*, in which multiplication of rDNA loci occurred, and compared them to three control species, namely *P. xylostella*, *S. frugiperda* and *L. dispar*, with a single major rDNA locus. In addition, *H. humuli* was compared to another hepialid, *P. californicus*. Combining molecular cytogenetics and sequencing of both second and third generation, we explored co-localization of 18S and 28S rDNA, association of major rDNA units with repetitive sequences, and their potential influence on evolution of major rDNA.

To determine the position and number of rDNA clusters, only one probe, often 18S rDNA, is hybridized on chromosomes as a representative of the whole major rDNA unit. It is usually assumed that major rDNA units spread as a whole (Bueno et al., 2013). However, Ferretti et al., (2019) discovered high intraspecific and interspecific variability and independent mobility of each component of the major rDNA unit (18S, ITS1, 5.8S, ITS2, and 28S) in the genome of six different populations of a grasshopper *Abracris flavolineata*. Similar pattern was also observed in *Coregonus* fishes where both complete and partial rDNA units were detected by FISH mapping of individual rRNA genes (Symonová et al., 2013). Therefore, we physically mapped partial sequences of 18S and 28S rDNA to test whether rDNA spreads as a whole unit in the studied species. In *H. humuli* and *C. ohridella*, both 18S and 28S rDNA signals co-localized and covered significant portion of one chromosome pair (Figure 1a, b) as reported earlier (Provazníková et al., 2021). Seven and eleven rDNA loci were highlighted by both probes in *A. urticae* and *I. io*, respectively (Figure 1c, d), which is in agreement with known distribution of their major rDNA (Nguyen et al., 2010; Provazníková et al., 2021). Thus, it is reasonable to conclude that a complete major rDNA unit was amplified or spread into new loci in the species under study. This is further corroborated by coverage analyses carried out in *H. humuli, A. urticae* and *I. io*, which showed similar read depth across their rDNA units (Figure 3 a, b, c, Suppl. Table S3).

For its dynamic evolution, rDNA has been often compared to repetitive sequences as arrays of rDNA are often found within heterochromatin (Lohe and Roberts, 2000). Fragments of rDNA can be amplified into satellite-like tandem arrays (Lohe and Roberts, 2000) and were found to be associated with satellites and other repeats (Barbosa et al., 2015; Jakubczak et al., 1991; Raskina et al., 2008; Sember et al., 2018; Symonová et al., 2013) which could mediate their spread (cf. Raskina et al., 2008; Nguyen et al., 2010; Proux-Wéra et al., 2013). Therefore, we clustered paired-end illumina reads of species under study using the RE pipeline (Novák et al., 2010) and searched for association of identified repeats with major rRNA genes. The paired-end reads did not link major rDNA with any other clustered repeats in *C. ohridella* and *A. urticae*,although we could not have excluded that such repeats are present either in low frequencies or distance bigger than the library insert size (see below). Yet, the association was recovered between major rDNA and Ty3/gypsy retrotransposon in *H. humuli* (Hh Ty3/gypsyA), and R2 element (IiR2) and a satellite (IiSat) in *I. io* (Suppl. Table S1).

Third generation sequencing technologies have recently provided and unprecedented insight into organisation of repetitive sequences including rDNA genes (e.g. Belser et al., 2021; Sims et al., 2021; Sproul et al., 2020; Vondrak et al., 2021). We took advantage of *A. urticae* and *I. io* PacBio data recently released by the Darwin Tree of Life project and analysed long reads, which contained major rDNA. The *I. io* data confirmed our previous results as the *I. io* long reads contained both the IiR2 and IiSat sequences (Suppl. Figure S3, Figure 4). Both proportion of long reads (Suppl. Figure S3) and coverage analysis (Figure 3c, Suppl. Table S3) suggest that IiR2 is present in about one third of rDNA units, which is in agreement with findings from other species (cf. Jakubczak et al., 1991; Zhou et al., 2013). In accordance with the results of RE, we found no MEs associated with rDNA in vast majority of reads in *A. urticae*. Small fraction of reads, which contained retrotransposon sequences of Ty3/Gypsy, R1 and R2 (Suppl. Figure S4), most likely corresponds to pseudogenes resulting from birth-and-death process (cf. Martí et al., 2021). Yet in contrast to the RE analysis, we found also satellite arrays (AuSat) in the IGS regions in majority of the *A. urticae* long reads bearing major rDNA (Suppl. Figure S4, Figure 4). Surprisingly, different satellites are associated with major rDNA in the two closely related nymphalids. In both species, spacers notably varied in their length which points to a possible lack of concerted evolution. We hypothesize this is due to high number of major rDNA loci, which are not all transcriptionally active. Thus, they do not associate in nucleolus and evolve independently. Alternatively, the observed variation could be ascribed to rDNA subtypes with tissue specific expression or to mutations impairing chromatin modification enzymes (cf. Havlová et al., 2016).

**Figure 4:**
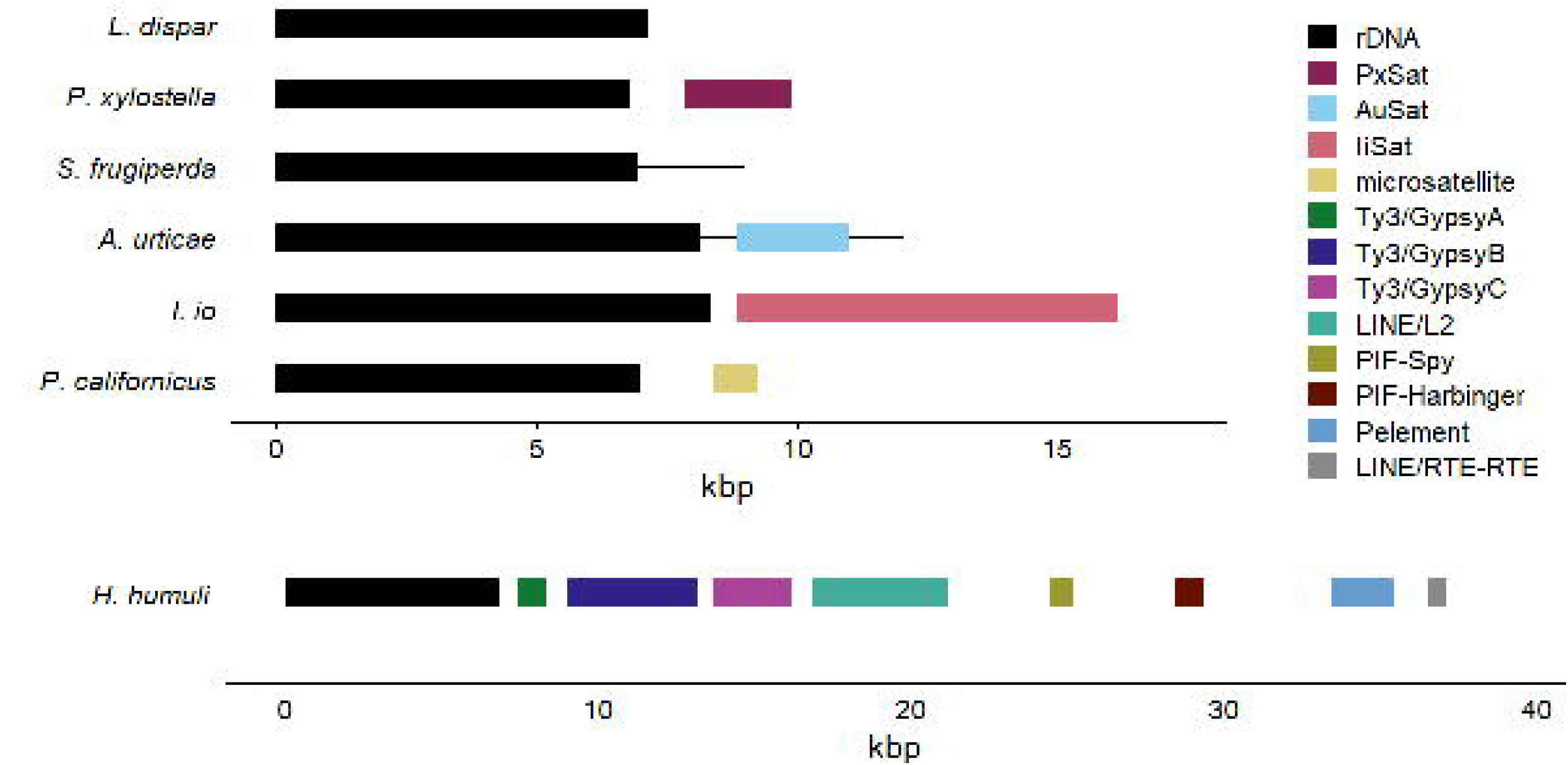
Schematic representation of most observed rDNA unit based on long read analysis. Black line represents the length of rDNA unit and coloured blocks position of rDNA genes and the repetitive elements.

In *H. humuli*, the Hh Ty3/gypsyA retrotransposon was inserted at the very end of rDNA unit, at the junction between 28S rDNA gene and external transcribed spacer (Suppl. Figure 1, Figure 4). Mapping of its partial sequence by TSA FISH revealed its clear co-localization with rDNA (Figure 2a-d). Although we did not detect any other Hh Ty3/gypsyA loci, we cannot exclude it is present elsewhere in the genome as interspersed repeats as combined length of used probes is still under ca. 1300 bp detection limit of the TSA FISH protocol used (cf. (Carabajal Paladino et al., 2014). Indeed, the coverage analysis suggests that abundance of MEs associated with rDNA is higher than abundance of rRNA genes themselves (Figure 3a, Suppl. Table S3). Furthermore, the Hh Ty3/gypsyA copy associated with major rDNA is non-autonomous. It represents only a portion of the corresponding RE cluster, which can be assembled into a complete retrotransposon with LTR repeats. Yet the Hh Ty3/gypsyA sequences in IGS lack long terminal repeats and most of protein coding domains (Suppl. Figure 1, Figure 4). Similar association between major rDNA and the Ty3/gypsy retrotransposon *Beon1 (* Galadriel clade) was observed also in the beet *Beta vulgaris* where, however, the ME is inserted into 18S rRNA genes (Weber et al. 2013).

While short paired-end reads revealed only association between rDNA and Hh Ty3/gypsyA retrotransposon in *H. humuli*, long ONT reads showed that fragments of eight different elements were inserted in IGS (Suppl. Figure 1, Figure 4). Like the Hh Ty3/gypsyA (see above), none of the major rDNA associated repeats is autonomous and thus cannot multiply on their own. However, their transmission is ensured by hitchhiking along with the indispensable rRNA gene family. It is not clear how expansion of IGS effected transcription of major rRNA genes. The IGS expanded to ca. 39 kb and it is roughly on par with 45 kb long IGS of mouse (Grozdanov et al., 2003) and thus not necessarily detrimental. If the IGS expansion decreases expression of major rRNA genes, increase of copy number would be favoured, which would explain the extraordinary size of the *H. humuli* major rDNA cluster. Moreover, it is possible that rDNA did not actually spread along the chromosome. Rather the total size of the array could have increased as major rDNA unit expanded due to insertion of repeats into IGS. Comparison of rDNA sequences between *H. humuli* and another hepialid *P. californicus*,showed that the expansion of IGS is not shared across the family Hepialidae. *P. californicus* IGS and major rDNA transcription unit have in total only ca. 10 kb. Surprisingly, the IGS contains an array of insect telomeric motif TTAGG_(n)_ and TTATG microsatellite (Suppl. Figure 2, Figure 4; cf. Ruiz-Herrera et al., 2008; Scali et al., 2016; Sember et al., 2018). IGSs are known to contain repetitive motives, which usually do not correspond to MEs (Grozdanov et al., 2003; Havlová et al., 2016). However, association between rDNA and MEs has been reported with 5S being involved more often than 45S rDNA (da Silva et al., 2016; Yano et al., 2020). Insertions of MEs into IGS observed in *H. humuli* thus represents interesting case of complex repeat organization (cf. Vondrak et al., 2021).

To pinpoint a mechanism responsible for changes in distribution of major rDNA in Lepidoptera, we took advantage of available long read sequencing data and compared structure of major rDNA units between species with the highly derived rDNA distribution (see above) with three species with a single rDNA locus, namely *P. xylostella*, *L. dispar*, and *S. frugiperda* (Nguyen et al., 2010; Provazníková et al., 2021). While we found MEs and/or satellite arrays at least in part of the IGS in all species with extraordinary major rDNA patterns but *C. ohridella* for which long reads have not been available, no repetitive sequences were associated with major rDNA in *S. frugiperda* (Suppl. Figure S7, Figure 4). However, satellite arrays (PxSat) of variable size were found in IGS region in *P. xylostella* (Suppl. Figure S5, Figure 4) and analysis of *L. dispar* long reads revealed two types of R1 retrotransposon (Suppl. Figure S6) with 52.3 % nucleotide identity in small fraction of reads.

The R1 and R2 non-LTR retrotransposons are, along with the Pokey DNA transposon, among the few known rDNA-specific elements with insertion sites in the gene for 28S rRNA (Eickbush and Eickbush, 2007, Elliott et al., 2013). The R elements represent one of the oldest groups of metazoan MEs (Kojima and Fujiwara, 2005). They were described for the first time in a fruit fly, *D. melanogaster*, by Roiha et al., (1981) and afterwards reported in many other animal phyla (reviewed in Eickbush and Eickbush, 2015). Within the order Lepidoptera, the R1 and/or R2 elements have so far been detected only in several species, namely *Bombyx mori* and *B. mandarina* (both Bombycidae), *Manduca sexta* (Sphingidae), four representatives of the family Saturniidae, and *L. dispar* (Erebidae) (Fujiwara et al., 1984; Jakubczak et al., 1991; Kierzek et al., 2009). The latter is of particular interest as our observation of the two R1 retrotransposons and no R2 element in the same species suggests interpopulation differences and rapid turnover of the R1 and R2 retrotransposons. This is further supported by the absence of the R2 element in the Czech *A. urticae* compared to the British population. The R elements were found also in the nymphalids *A. urticae* and *I. io* with seven and eleven rDNA loci, respectively. It was proposed that the R2 retrotransposon plays an important role in maintaining rDNA copy numbers in *Drosophila* (Nelson et al., 2021). Yet, their absence in *S. frugiperda* and *P. xylostella* does not destabilize their rDNA loci and it seems unlikely that the R1 and R2 retrotransposons could mobilize rDNA in species under study. Although it was shown that the R1 and R2 retrotransposons can insert in target site outside 28S rDNA in *B. mori* (Xiong et al., 1988), the IiR2 elements were not detected outside the major rDNA loci in *I. io* (Figure 2b, 3b). Moreover, insertion of the R1 and R2 elements into the 28S rDNA cause pseudogenization of the corresponding rDNA units (Long and Dawid, 1979). The only other MEs associated with major rDNA were observed in *H. humuli*. However, these were not autonomous and their sequence diverged from those found in the rest of the genome. Thus, it seems unlikely that MEs could mediate spread of rDNA observed in some Lepidoptera (Provazníková et al., 2021) via transposition.

On the contrary, satellite arrays such as those found in IGS of nymphalids with high number of rDNA loci, *A. urticae* and *I. io* (Figure 4), could facilitate homology-mediated spread of rDNA via either ectopic recombination or integration of extrachromosomal rDNA circles (Muirhead and Presgraves, 2021; Nguyen et al., 2010; Proux-Wéra et al., 2013; Sproul et al., 2020). Yet, satellite arrays were associated with rDNA also in *P. xylostella* (Figure 4), which has only one rDNA locus. There is a difference between satellite DNA associated with rDNA in *P. xylostella* and the two nymphalids. In *P. xylostella*, we did not find any long reads bearing the PxSat without rDNA or at least partial IGS sequence. In both nymphalids, however, we identified more than 10 long reads bearing only the satellite arrays, with mean IGS and filtered read lengths being similar in all species which points to a presence of the satellites outside rDNA arrays. Unfortunately, coverage analyses are uninformative for satellites as there is a considerable variation in number of their monomers per rDNA unit (Suppl. Figure S5). Inspection of the *A. urticae* genome assembly suggests that the AuSat is localized only in rDNA clusters and arrays adjacent to rDNA (Suppl. Figure S4), which suggests that rDNA spread into the AuSat loci.

We cannot distinguish with certainty whether the rDNA spread occurred via ectopic recombination or integration of ecc-rDNA. Yet we argue that preferential spread of rDNA into terminal regions of lepidopteran chromosomes (Nguyen et al., 2010; Provazníková et al., 2021) favours ectopic recombination as its efficiency depends on proximity of homologous sequences to telomeres (Goldman and Lichten, 1996; Nguyen et al., 2010), whereas ecc-rDNA could be integrated anywhere in the genome as long as a homologous sequence is present. Little is known about satellite DNA in Lepidoptera, which has been studied in detail only in a dozen of species (Lu et al., 1994; Mandrioli et al., 2003; Mahendran et al., 2006; Věchtová et al., 2016; M. Dalíková et al., 2017b; Cabral-de-Mello et al., 2021). Yet, it seems that abundance of satellite DNA in lepidopteran genomes is very low with scattered distribution and possible enrichment on sex chromosomes (Cabral-de-Mello et al., 2021). This does not reflect distribution of rDNA in Lepidoptera (Nguyen et al., 2010; Provazníková et al., 2021). Yet we cannot exclude those satellites associated with major rDNA are limited to chromosome ends similar to *P. californicus*, which contains telomeric repeats in its IGS (Suppl. Figure S2).

Compared to other regions of rDNA arrays, IGS are rarely studied, and we thus cannot tell whether the observed complex association between rDNA and repetitive sequences (Figure 4) represent a common phenomenon. Our results show that long read sequencing is a valuable tool to study association of repeats including major rDNA as it provided more detailed information about major rDNA associated repeats than analysis of short reads limited by library insert size. Moreover, the long read analysis provided better genomic representation compared to the genome assembly based on these long reads as seen in the *A. urticae* example (Suppl. Figure 4 and 9). Available target enrichment of major rDNA and other repeats for long read sequencing (McKinlay et al., 2021) could provide further insight into formation of complex repeat structures involving rDNA.

## Material and methods

### Material

Specimens of all studied species were collected from wild populations. Females of *Hepialus humuli* were collected in Bochov, Czech Republic and let lay eggs in plastic containers. Hatched larvae were transferred and extensively reared in outdoor pots with planted carrot (*Daucus carota*). Larvae of *Cameraria ohridella* were collected in České Budějovice, Czech Republic, from leaves of the horse-chestnut, *Aesculus hippocastanum*. Specimens of *C. ohridella* were processed immediately after collection. Larvae of two nymphalids, *Inachis io* and *Aglais urticae*, were collected near Vrábče, Czech Republic. They were kept on the common nettle (*Urtica dioica*) in ambient conditions. Larvae of *Phymatopus californicus* were collected from the yellow bush lupine, *Lupinus arboreus*, in the Bodega Marine Reserve (California, USA). Larvae were used for chromosomal preparations and extraction of genomic DNA (gDNA) shortly after collection.

### Chromosomal preparation

Chromosomal preparations were prepared by a spreading technique as described in Mediouni et al., (2004) and Dalíková et al., (2017a) with 10 min hypotonization of tissue. Meiotic and mitotic preparations were obtained from gonads of late larval instars of all four species. Afterwards, prepared slides were dehydrated in an ethanol series (70, 80 and 100% ethanol, 30 sec each) and stored at −20 and −80 °C until further use. Remaining tissues were frozen for subsequent gDNA extraction.

### Genomic DNA extraction

For downstream applications such as PCR, gDNA was extracted from larvae using NucleoSpin DNA Insect (Macherey-Nagel, Düren, Germany) according to manufacturer’s protocol. To obtain high-molecular-weight gDNA for NGS sequencing, gDNA was extracted from larvae using MagAttract HMW DNA Kit (Qiagen, Hilden, Germany) or Nanobind Tissue Big DNA kit (Circulomics Inc, Baltimore, MD, USA) according to manufacturer’s protocol. Concentration of extracted samples was measured by Qubit 3.0 Fluorometer (Invitrogen, Carlsbad, CA, USA) and visualized on agarose gel. A single male larva was used as input material for all species but *C. ohridella*, for which 5-10 individuals (larvae and pupae) of both sexes were pooled because of their small size. A single male adult was used for extraction for Nanopore sequencing.

### Repeat Explorer analysis

For analysis of repetitive DNA content, whole gDNA was sequenced on the Illumina platform generating either 150 bp pair-end reads from library with mean insert size 450 bp (Novogene Co., Ltd., Beijing, China) or 250 bp PE reads with the mean insert size 700 bp in case of *C. ohridella* (Genomics Core Facility, EMBL Heidelberg, Germany). The raw reads were quality filtered and trimmed to uniform length of 120 bp (230 bp for *C. ohridella*) by Trimmomatic 3.2 (Bolger et al., 2014). Random sample of two million (one million for *C. ohridella*) trimmed PE reads was analysed by RE pipeline (version cerit-v0.3.1-2706) implemented in Galaxy environment (https://repeatexplorer-elixir.cerit-sc.cz/galaxy/) with automatic annotation via blastn and blastx using the Metazoan 3 Repeat Explorer database. The resulting html files were searched for clusters annotated as major rDNA and their connection to other clusters.

### Probes for FISH experiment

All mapped sequences were amplified by PCR using specific primers (for details see in Table 1), purified from agarose gel, and cloned into Promega pGem T-Easy Vector (Promega, Madison, WI, USA). Selected clones were isolated using Nucleo Spin Plasmid kit (Macherey-Nagel) and verified by sequencing (SEQme, Dobříš, Czech Republic).

**Table 1:**
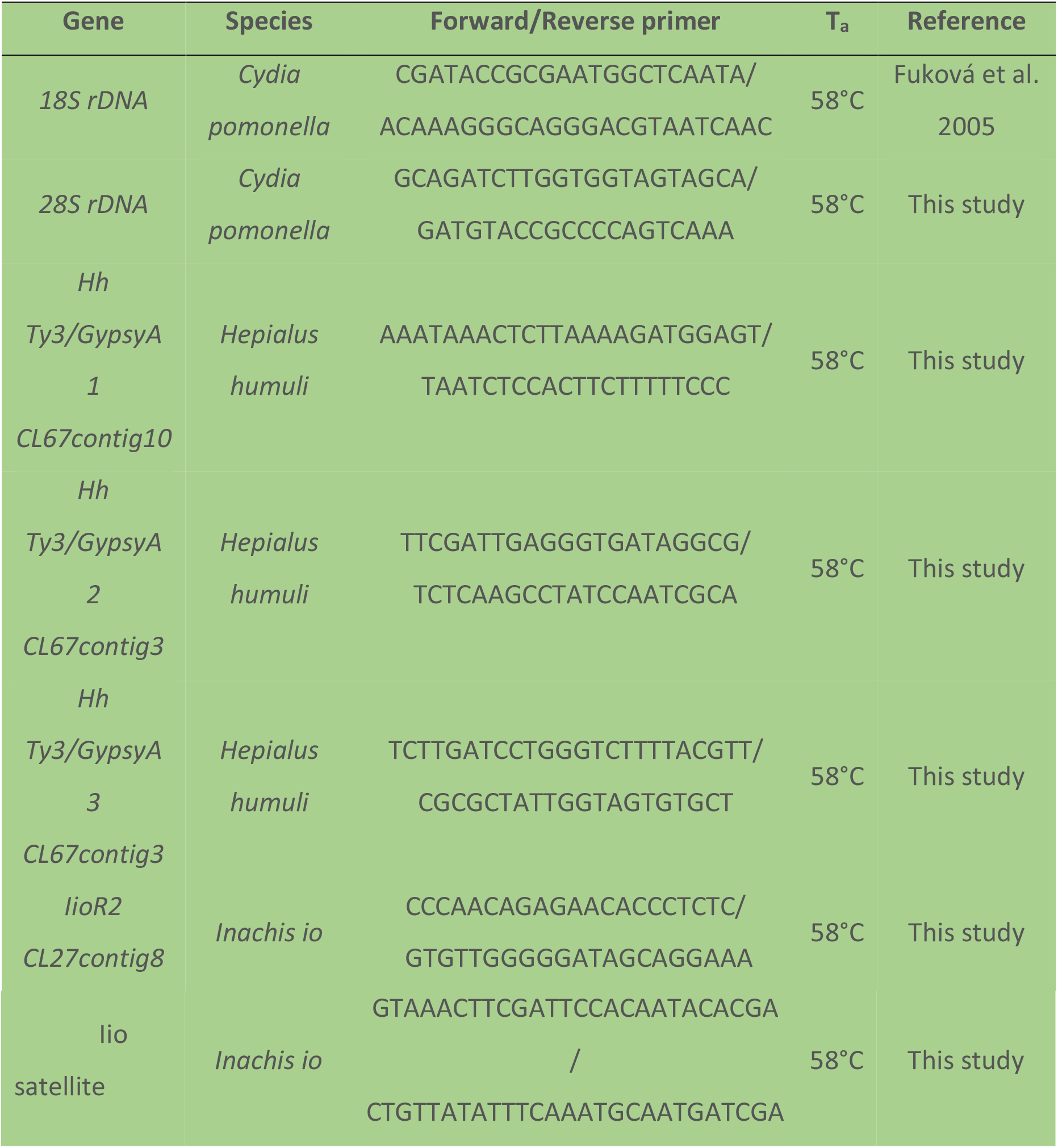
Primers used for PCR amplification of FISH probe templates and labelling.

Fragments of 18S and 28S rRNA genes were amplified from gDNA of codling moth, *Cydia pomonella* (Tortricidae) (cf. Nguyen et al., 2010). To obtain probes, these fragments were re-amplified by PCR from plasmids using specific primers (Table 1), purified by Wizard^®^ SV Gel and PCR Clean-Up System (Promega), and labelled using nick translation protocol by (Kato et al., 2006) with modifications described in (Martina Dalíková et al., 2017a). The 20μL labelling reaction contained 1 μg DNA; 0.5 mM each dATP, dCTP, and dGTP; 0.1 mM dTTP; 20 μM labelled nucleotides; 1× nick translation buffer (50 mM Tris–HCl, pH 7.5; 5 mM MgCl2; 0.005% BSA); 10 mM β-mercaptoethanol; 2.5 × 10^-4^ U DNase I, and 1 U DNA polymerase I (both ThermoFisher Scientific, Waltham, MA). The reaction was incubated at 15 °C for 45 minutes and enzymes were inactivated at 70 °C for 10 minutes. The 18S rDNA probe was labelled either with biotin-16-dUTP (Roche Diagnostics, Basel, Switzerland) or DNP-11-dUTP (Jena Bioscience, Jena, Germany) and 28S rDNA probe was labelled by digoxigenin-11-dUTP (Roche Diagnostics) or fluorescein-12-dUTP (Jena Bioscience).

Three sequence fragments of Hh Ty3/GypsyA (CL67 contig 3 and 10) found in *H. humuli* and one of R2 element (CL27contig8) and satellite (CL37 contig4) found in *I. io*, were separately labelled by PCR using plasmid DNA as template according to Provazníková et al., (2021). The 25μL labelling reaction contained 1 – 10 ng template plasmid DNA, 1x Ex Taq buffer, 1 mM each dATP, dCTP, and dGTP; 0.36 mM dTTP; 0.62 mM labelled nucleotides of Fluorescein-12-ddUTP (Jena Biosciences), 5 μmol of each specific primer (**Table 1**), and 0.25 U TaKaRa Ex Taq DNA polymerase (TaKaRa, Otsu, Japan). The resulting labelled probes were purified using Sephadex gel filtration (Illustra Sephadex G-50 fine DNA grade).

### FISH with 18S and 28S rDNA probes

Indirect FISH was carried out according to Fuková et al., (2005) and Zrzavá et al., (2018) with some modifications and using two probes, biotin-labelled 18S rDNA probe and digoxigenin-labelled 28S rDNA probe, simultaneously. This technique was used to localize 18S and 28S rDNA genes in genome of *H. humuli*, *C. ohridella*, and *A. urticae*. The slides were dehydrated in an ethanol series (70, 80, and 100% ethanol, 30 sec each) and pre-treated with RNase A (200 ng/μL) in 2× SSC at 37°C for 1 h, washed twice in 2×SSC at RT for 5 min each and incubated in 5×Denhardt’s solution at 37°C for 30 minutes. After, slides were denaturated in 70% formamide in 2x SSC at 68°C for 3.5 minutes and immediately dehydrated in an ethanol series (cold 70% for 1 min, 80 and 100% for 30 sec each). Hybridization probe mix containing 10% dextran sulfate, 50% deionized formamide, 25 μg of sonicated salmon sperm and 50 ng of each probe in 2× SSC in final volume of 10 μl was denaturated at 90°C for 5 minutes and immediately placed on ice for 2 minutes. Afterwards, hybridization probe mix was applied on the slide, covered by cover slip, and placed into a humid chamber. Hybridization was carried out at 37°C overnight (12-16 h).

Next day, slides were incubated three times in 50% formamide in 2x SSC and followed by three washes in 2x SSC, both at 46°C for 5 minutes. Slides were then washed three times with 0.1x SSC at 62°C for 5 minutes and once in 1% Triton X in 4x SSC at RT for 10 minutes. The slides were blocked with 2.5% BSA in 4x SSC at RT for 30 minutes, incubated with anti-DIG1 (mouse anti-digoxigenin, 1:100, Roche Diagnostics, Basel, Switzerland) and streptavidin Cy3 conjugate (1:1000, Jackson ImmunoRes. Labs. Inc, West Grove, PA, USA) in 2.5% BSA in 4x SSC at 37°C for 1 hour and washed three times with 1% Triton X in 4x SSC at 37°C for 3 minutes each. To amplify the signals, last three steps were repeated twice, firstly with anti-DIG2 (sheep anti-mouse Ig digoxigenin conjugate, 1:200, Merck Millipore, Billerica, MA, USA) and antistreptavidin (1:25, Vector Labs. Inc, Burlingame, CA, USA) and secondly with anti-DIG3 (sheep anti-digoxigenin fluorescein conjugate, 1:200, Roche Diagnostics) and streptavidin-Cy3 conjugate (1:1000, Jackson ImmunoRes. Labs. Inc). After the last washing step, slides were incubated in 1% Kodak PhotoFlo at RT for 1 minute and counterstained with 0.5 mg/mL DAPI (4′,6-diamidino-2-phenylindole, Sigma-Aldrich, St. Louis, MO, USA) in antifade based on DABCO (1,4-diazabicyclo(2.2.2)-octane; Sigma–Aldrich).

### Double TSA FISH

Double FISH with tyramide signal amplification (double TSA FISH) was performed according to Carabajal Paladino et al., (2014) with some modifications. Due to its high sensitivity, double TSA FISH was employed to localize 18S and 28S rDNA genes in *I. io* genome, and ME sequences and 18S rDNA gene in genomes of *I. io* and *H. humuli*. Briefly, slides were dehydrated in an ethanol series (70,80 and 100% ethanol, 30 sec each) and pre-treated with 50 μg/mL pepsin in 0.01 M HCl at 37°C for 10 min, 1% H_2_O_2_ in 1x PBS at RT for 30 min, and in RNase A (100 μg/mL) in 1x PBS at 37°C for 1 hour. After each pre-treatment, the slides were washed three times in 1x PBS at RT for 5 min each washing. After the last washing, slides were incubated in 5x Denhardt’s solution at 37°C for 30 minutes. Directly after the last incubation, 50 μl of hybridization probe mix containing 10% dextran sulphate, 50% deionized formamide, and 10-20 ng of each probe in 2x SSC was applied onto the slide, covered by cover slip, and incubated at 70°C for 5 minutes. Afterwards, slides were placed into the humid chamber and hybridized at 37°C overnight (12-16 h). In *I. io* experiments, the 18S rDNA probe (10-20 ng) labelled with dinitrophenyl (DNP) was used with fluorescein-labelled R2 probe (10-20 ng), or fluorescein-labelled 28S rDNA probe (10-20 ng). In case of *H. humuli*, combination of three fluorescein-labelled ME probes (Hh Ty3/GypsyA 1-3, 5-10 ng each) and 18S rDNA probe (10-20 ng) was used.

The next day, slides were incubated three times 5 min in 50% formamide in 2x SSC at 46°C each, washed three times in 2x SSC at 46°C for 5 minutes each and in 0.1x SSC at 62°C for 5 minutes each, and washed once in 1x TNT at RT for 5 minutes. The slides were blocked in TNB buffer at RT for 30 minutes and incubated with antifluorescein-HRP Conjugate (PerkinElmer) in TNB (diluted 1:1000) at RT for 1 hour. Afterwards, the slides were washed three times in 1x TNT at RT for 5 minutes each and incubated with TSA Plus Fluorescein (PerkinElmer) according to the manual at RT for 3-15 minutes (3-5 min in *H. humuli* and *C. ohridella*, 10-15 min for *I.io*) and washed again three times in 1x TNT at RT for 5 minutes each. To perform the second round of detection and to quench peroxidase activity from previous steps, slides were incubated in 1% H_2_O_2_ in 1x PBS at RT for 30 min. Next, the slides were washed three times in 1x TNT at RT for 5 minutes each and the amplification steps were repeated using anti-DNP-HRP Conjugate (PerkinElmer) and TSA Plus Cyanine 3 (PerkinElmer). After the last washing step, the slides were incubated in 1% Kodak PhotoFlo at RT for 1 minute and counterstained with 0.5 mg/mL DAPI (Sigma–Aldrich) in antifade DABCO (Sigma–Aldrich).

### Microscoping and image processing

Chromosome preparations from FISH experiments were observed by Zeiss Axioplan 2 microscope (Carl Zeiss, Jena, Germany) equipped with appropriate fluorescence filter sets. An Olympus CCD monochrome camera XM10 equipped with cellSens 1.9 digital imaging software (Olympus Europa Holding, Hamburg, Germany) was used to record and capture black-and-white pictures. Images were captured separately for each fluorescent dye and then pseudocoloured and superimposed with Adobe Photoshop CS4, version 11.0.

### Long read sequencing and analysis

High molecular weight DNA from *H. humuli* was enriched for fragments longer than 10 kbp by Short Read Eliminator (Circulomics Inc). The library was prepared by Ligation Sequencing Kit SQK-LSK110 (Oxford Nanopore Technologies, Oxford, UK) according to the manufacture’s protocol and therein recommended third party consumables. The library was snap-frozen and stored over night at −70°C and then sequenced using flowcell R10.3 and MinION Mk1B (Oxford Nanopore Technologies). Reads were basecalled by guppy 4.4.1. with high accuracy flip-flop algorithm. The data was filtered for reads 15kbp and longer with quality score over 10 using NanoFilt (De Coster et al., 2018).

Quality and length filtered reads were searched for presence of major rDNA using blastn. Reads containing at least 1000 bp of *H. humuli* major rDNA unit were assembled by Flye 2.8 (Kolmogorov et al., 2019) using minimal overlap 8 kbp. The annotation of MEs was done by RepeatMasker 4.1.2-p1 (Smit et al., 2013) protein-based masking. Tandem repeats were identified based on self Dotplot implemented in Geneious 11.1.5. Consensus sequences of all identified ME fragments together with major rDNA unit were mapped to individual rDNA bearing nanopore reads using minimap2 (Li, 2018) with appropriate pre-set. The presence and relative localization of individual elements was evaluated via R script (R version 4.0.3 in Rstudio version 1.4.1103). Only regions with mapping quality at least 20 were considered.

*Phymatopus californicus* gDNA was sequenced on Oxford Nanopore platform in Novogene Co.,Ltd. PacBio HiFi reads of *I. io* (project PRJEB42130) and *A. urticae* (project PRJEB42112) were obtain through the Darwin Tree of Life project (http://www.darwintreeoflife.org). PacBio CLR data were obtained from Sequence Read Archive (SRA) database (*S. frugiperda* SRR12642577; *L. dispar* SRR13505170-6, SRR13505182-3, and SRR13505187; *P.xylostella* SRR13530960). Further, the reads were processed same as in *H. humuli* except for the HiFi reads, which were not quality filtered.

Similar approach to detect rDNA and associated repetitive DNA was used also in *A. urticae* chromosomal level genome assembly (Bishop et al., 2021) (ENA acc. No. PRJEB41896).

### Coverage analysis

Coverage analysis was done by aligning genomic Illumina sequencing reads from *H. humuli, I. io*, and *A. urticae* to consensus sequences, which were generated by overlapping the contigs from RE in Geneious 11.1.5 or by Flye 2.8 assembler, using Bowtie2 aligner (Langmead et al., 2019; Langmead and Salzberg, 2012). Coverage values were obtained using samtools depth (v 1.10) (Li et al., 2009) and plotted using a script in R (R version 4.1.0 in Rstudio Workbench Version 1.4.1717-3). Mean coverage of defined annotation blocks as seen in Figure 3 was computed using R and is in Suppl. Tables 3.

## Data availability

Sequencing data generated in this study was deposited in the NCBI Sequence read archive under Bioproject reg. no. PRJNA737195. Long reads bearing rDNA of species under study, assemblies of their rDNA units and R codes used for analyses were deposited in the Dryad Digital Repository under doi: 10.5061/dryad.gmsbcc2qj.

## Acknowledgement

We would like to thank our colleagues Anna Voleníková, František Marec, Leonela Carabajal-Paladino, Magda Zrzavá, Vladimír Beneš, and Marie Korchová for their help and extensive support. We would also like to show our gratitude to the Darwin Tree of Life project for allowing us to use its HiFi PacBio data. Computational resources were provided by the CESNET LM2015042 and the CERIT Scientific Cloud LM2015085, provided under the programme “Projects of Large Research, Development, and Innovations Infrastructures”. Computational resources were also provided by the ELIXIR-CZ project (LM2015047), part of the international ELIXIR infrastructure.

## Supplemental material legends

**Figure S1:** Visualization of major rDNA genes and associated repeats on *H. humuli* Oxford Nanopore reads. Only matches longer than 200 bp and with mapping quality over 20 were considered. Gray line represents the read length, coloured boxes correspond to the rDNA genes or individual detected repeats.

**Figure S2:** Visualization of major rDNA genes and associated repeats on *P. californicus* Oxford Nanopore reads. Only matches longer than 200 bp and with mapping quality over 20 were considered. Gray line represents the read length, coloured boxes correspond to the rDNA genes or individual detected repeats.

**Figure S3:** Visualization of major rDNA genes and associated repeats on *I. io* HiFi PacBio reads. Only matches longer than 200 bp and with mapping quality over 20 were considered. Gray line represents the read length, coloured boxes correspond to the rDNA genes or individual detected repeats.

**Figure S4:** Visualization of major rDNA genes and associated repeats on *A. urticae* HiFi PacBio reads. Only matches longer than 200 bp and with mapping quality over 20 were considered. Gray line represents the read length, coloured boxes correspond to the rDNA genes or individual detected repeats.

**Figure S5:** Visualization of major rDNA genes and associated satellite on *P. xylostella* PacBio reads. Only matches longer than 200 bp and with mapping quality over 20 were considered. Gray line represents the read length, coloured boxes correspond to the rDNA genes or PxSat.

**Figure S6:** Visualization of major rDNA genes and associated repeats on *L. dispar* PacBio reads. Only matches longer than 200 bp and with mapping quality over 20 were considered. Gray line represents the read length, coloured boxes correspond to the rDNA genes or individual detected repeats.

**Figure S7:** Visualization of major rDNA genes *S. frugiperda* PacBio reads. Only matches longer than 200 bp and with mapping quality over 20 were considered, no associated repeats were detected. Gray line represents the read length, coloured boxes correspond to the rDNA genes.

**Figure S8:** Distribution of AuSat (in *A. urticae*), IiSat (in *I. io*), and PxSat (in *P. xylostella*) arrays lengths in PacBio reads containing at least 500bp of major rDNA genes. Bold line represents median value, box corresponds to interquartile range, and the outlier values are represented by individual dots.

**Figure S9:** Visualization of major rDNA genes and associated repeats from the current *A. urticae* genome assembly. Only matches longer than 200 bp and with mapping quality over 20 were considered. Gray line represents the chromosomal length scaffolds with detected rDNA clusters, coloured boxes correspond to the rDNA genes or individual detected repeats.

**Table S1:** Summary of basic characteristics of selected clusters from Repeat Explorer analysis in *H. humuli*, *A. urticae*, *I. io*, and *C. ohridella*

**Table S2:** Length characterisation of PacBio reads and satellite arrays in *A. urticae*, *I. io*, and *P. xylostella*

**Table S3:** Summary of coverage analysis of individual elements in rDNA units in *H. humuli*, *A. urticae*, and *I. io*

## References

Ahola V, Lehtonen R, Somervuo P, Salmela L, Koskinen P, Rastas P, Välimäki N, Paulin L, Kvist J, Wahlberg N, Tanskanen J, Hornett EA, Ferguson LC, Luo S, Cao Z, de Jong MA, Duplouy A, Smolander O-P, Vogel H, McCoy RC, Qian K, Chong WS, Zhang Q, Ahmad F, Haukka JK, Joshi A, Salojärvi J, Wheat CW, Grosse-Wilde E, Hughes D, Katainen R, Pitkänen E, Ylinen J, Waterhouse RM, Turunen M, Vähärautio A, Ojanen SP, Schulman AH, Taipale M, Lawson D, Ukkonen E, Mäkinen V, Goldsmith MR, Holm L, Auvinen P, Frilander MJ, Hanski I. 2014. The Glanville fritillary genome retains an ancient karyotype and reveals selective chromosomal fusions in Lepidoptera. Nat Commun 5:4737. doi:10.1038/ncomms5737

Barbosa P, de Oliveira LA, Pucci MB, Santos MH, Moreira-Filho O, Vicari MR, Nogaroto V, de Almeida MC, Artoni RF. 2015. Identification and chromosome mapping of repetitive elements in the *Astyanax scabripinnis* (Teleostei: Characidae) species complex. Genetica 143:55–62. doi:10.1007/s10709-014-9813-2

Baumgärtner L, Paiz LM, Zawadzki CH, Margarido VP, Portela Castro ALB. 2014. Heterochromatin polymorphism and physical mapping of 5S and 18S ribosomal DNA in four populations of *Hypostomus strigaticeps* (Regan, 1907) from the Paraná river basin, Brazil: Evolutionary and environmental correlation. Zebrafish 11:479–487. doi:10.1089/zeb.2014.1028

Bedo DG. 1984. Karyotypic and chromosome banding studies of the potato tuber moth, *Phthorimaea operculella* (Zeller) (Lepidoptera, Gelechiidae). Can J Genet Cytol 26:141–145. doi:10.1139/g84-024

Belser C, Baurens F-C, Noel B, Martin G, Cruaud C, Istace B, Yahiaoui N, Labadie K, Hřibová E, Doležel J, Lemainque A, Wincker P, D’Hont A, Aury J-M. 2021. Telomere-to-telomere gapless chromosomes of banana using nanopore sequencing. Commun Biol 4:1047. doi:10.1038/s42003-021-02559-3

Bishop G, Ebdon S, Lohse K, Vila R, Darwin Tree of Life Barcoding collective, Wellcome Sanger Institute Tree of Life programme, Wellcome Sanger Institute Scientific Operations: DNA Pipelines collective, Tree of Life Core Informatics collective, Darwin Tree of Life Consortium. 2021. The genome sequence of the small tortoiseshell butterfly, *Aglais urticae* (Linnaeus, 1758). Wellcome Open Res 6:233. doi:10.12688/wellcomeopenres.17197.1

Bolger AM, Lohse M, Usadel B. 2014. Trimmomatic: A flexible trimmer for Illumina sequence data. Bioinformatics 30:2114–2120. doi:10.1093/bioinformatics/btu170

Bueno D, Palacios-Gimenez OM, Cabral-de-Mello DC. 2013. Chromosomal mapping of repetitive DNAs in the grasshopper *Abracris flavolineata* reveal possible ancestry of the B chromosome and H3 histone spreading. PLoS ONE 8:e66532. doi:10.1371/journal.pone.0066532

Cabral-de-Mello DC, Oliveira SG, de Moura RC, Martins C. 2011. Chromosomal organization of the 18S and 5S rRNAs and histone H3 genes in Scarabaeinae coleopterans: insights into the evolutionary dynamics of multigene families and heterochromatin. BMC Genetics 12:88. doi:10.1186/1471-2156-12-88

Cabral-de-Mello DC, Zrzavá M, Kubíčková S, Rendón P, Marec F. 2021. The role of satellite DNAs in genome architecture and sex chromosome evolution in Crambidae moths. Front Genet 12:661417. doi:10.3389/fgene.2021.661417

Carabajal Paladino LZ, Nguyen P, Šíchová J, Marec F. 2014. Mapping of single-copy genes by TSA-FISH in the codling moth, *Cydia pomonella*. BMC Genet 15:S15. doi:10.1186/1471-2156-15-S2-S15

Carabajal Paladino LZ, Provazníková I, Berger M, Bass C, Aratchige NS, López SN, Marec F, Nguyen P. 2019. Sex chromosome turnover in moths of the diverse superfamily Gelechioidea. Genome Biol Evol 11:1307–1319. doi:10.1093/gbe/evz075

Cheng T, Wu J, Wu Y, Chilukuri RV, Huang L, Yamamoto K, Feng L, Li W, Chen Z, Guo H, Liu J, Li S, Wang X, Peng L, Liu D, Guo Y, Fu B, Li Z, Liu C, Chen Y, Tomar A, Hilliou F, Montagné N, Jacquin-Joly E, d’Alençon E, Seth RK, Bhatnagar RK, Jouraku A, Shiotsuki T, Kadono-Okuda K, Promboon A, Smagghe G, Arunkumar KP, Kishino H, Goldsmith MR, Feng Q, Xia Q, Mita K. 2017. Genomic adaptation to polyphagy and insecticides in a major East Asian noctuid pest. Nat Ecol Evol 1:1747–1756. doi:10.1038/s41559-017-0314-4

da Silva M, Barbosa P, Artoni RF, Feldberg E. 2016. Evolutionary dynamics of 5S rDNA and recurrent association of transposable elements in electric fish of the family Gymnotidae (Gymnotiformes): The case of *Gymnotus mamiraua*. Cytogenet Genome Res 149:297–303. doi:10.1159/000449431

Dalíková Martina, Zrzavá M, Hladová I, Nguyen P, Šonský I, Flegrová M, Kubíčková S, Voleníková A, Kawahara AY, Peters RS, Marec F. 2017a. New insights into the evolution of the W chromosome in Lepidoptera. J Hered 108:709–719. doi:10.1093/jhered/esx063

Dalíková M., Zrzavá M, Kubíčková S, Marec F. 2017b. W-enriched satellite sequence in the Indian meal moth, *Plodia interpunctella* (Lepidoptera, Pyralidae). Chromosome Res 25:241–252. doi:10.1007/s10577-017-9558-8

De Coster W, D’Hert S, Schultz DT, Cruts M, Van Broeckhoven C. 2018. NanoPack: Visualizing and processing long-read sequencing data. Bioinformatics 34:2666–2669. doi:10.1093/bioinformatics/bty149

de Sene VF, Pansonato-Alves JC, Ferreira DC, Utsunomia R, Oliveira C, Foresti F. 2015. Mapping of the retrotransposable elements *Rex1* and *Rex3* in chromosomes of *Eigenmannia* (Teleostei, Gymnotiformes, Sternopygidae). Cytogenet Genome Res 146:319–324. doi:10.1159/000441465

Eickbush TH, Eickbush DG. 2015. Integration, regulation, and long-term stability of R2 retrotransposons. Microbiol Spectr 3:2. doi:10.1128/microbiolspec.MDNA3-0011-2014

Eickbush TH, Eickbush DG. 2007. Finely orchestrated movements: Evolution of the ribosomal RNA genes. Genetics 175:477–485. doi:10.1534/genetics.107.071399

Elliott TA, Stage DE, Crease TJ, Eickbush TH. 2013. In and out of the rRNA genes: characterization of *Pokey* elements in the sequenced *Daphnia* genome. Mobile DNA 4:20. doi:10.1186/1759-8753-4-20

Ferretti ABSM, Ruiz-Ruano FJ, Milani D, Loreto V, Martí DA, Ramos E, Martins C, Cabral-de-Mello DC. 2019. How dynamic could be the 45S rDNA cistron? An intriguing variability in a grasshopper species revealed by integration of chromosomal and genomic data. Chromosoma 128:165–175. doi:10.1007/s00412-019-00706-8

Fiore-Donno AM, Kamono A, Meyer M, Schnittler M, Fukui M, Cavalier-Smith T. 2012. 18S rDNA phylogeny of *Lamproderma* and allied genera (Stemonitales, Myxomycetes, Amoebozoa). PLoS ONE 7:e35359. doi:10.1371/journal.pone.0035359

Fujiwara H, Ogura T, Takada N, Miyajima N, Ishikawa H, Maekawa H. 1984. Introns and their flanking sequences of *Bombyx mori* rDNA. Nucl Acids Res 12:6861–6869. doi:doi: 10.1093/nar/12.17.6861

Fuková I, Nguyen P, Marec F. 2005. Codling moth cytogenetics: Karyotype, chromosomal location of rDNA, and molecular differentiation of sex chromosomes. Genome 48:1083–1092. doi:10.1139/g05-063

García-Souto D, Pérez-García C, Kendall J, Pasantes J. 2016. Molecular cytogenetics in trough shells (Mactridae, Bivalvia): Divergent GC-rich heterochromatin content. Genes 7:47. doi:10.3390/genes7080047

Goldman ASH, Lichten M. 1996. The efficiency of meiotic recombination between dispersed sequences in *Saccharomyces cerevisiae* depends upon their chromosomal location. Genetics 144:43–55. doi: 10.1093/genetics/144.1.43

Grozdanov P, Georgiev O, Karagyozov L. 2003. Complete sequence of the 45-kb mouse ribosomal DNA repeat: analysis of the intergenic spacer. Genomics 82:637–643. doi:10.1016/S0888-7543(03)00199-X

Havlová K, Dvořáčková M, Peiro R, Abia D, Mozgová I, Vansáčová L, Gutierrez C, Fajkus J. 2016. Variation of 45S rDNA intergenic spacers in *Arabidopsis thaliana*. Plant Mol Biol 92:457–471. doi:10.1007/s11103-016-0524-1

Ingle J, Timmis JN, Sinclair J. 1975. The relationship between satellite deoxyribonucleic acid, ribosomal ribonucleic acid gene redundancy, and genome size in plants. Plant Physiol 55:496–501. doi:10.1104/pp.55.3.496

Jakubczak JL, Burke WD, Eickbush TH. 1991. Retrotransposable elements RI and R2 interrupt the rRNA genes of most insects. PNAS 88:3295–3299,. doi:10.1073/pnas.88.8.3295

Kato A, Albert P, Vega J, Kato A, Albert P, Vega J, Birchler J. 2006. Sensitive fluorescence *in situ* hybridization signal detection in maize using directly labeled probes produced by high concentration DNA polymerase nick translation. Biotechnic Histochem 81:71–78. doi:10.1080/10520290600643677

Kierzek E, Christensen SM, Eickbush TH, Kierzek R, Turner DH, Moss WN. 2009. Secondary structures for 5’ regions of R2 retrotransposon RNAs reveal a novel conserved pseudoknot and regions that evolve under different constraints. J Mol Biol 390:428–442. doi:10.1016/j.jmb.2009.04.048

Kobayashi T. 2011. Regulation of ribosomal RNA gene copy number and its role in modulating genome integrity and evolutionary adaptability in yeast. Cell Mol Life Sci 68:1395–1403. doi:10.1007/s00018-010-0613-2

Kojima KK, Fujiwara H. 2005. Long-term inheritance of the 28S rDNA-specific retrotransposon R2. Mol Biol Evol 22:2157–2165. doi:10.1093/molbev/msi210

Kolmogorov M, Yuan J, Lin Y. 2019. Assembly of long, error-prone reads using repeat graphs. Nat Biotechnol 37:540–546. doi:10.1038/s41587-019-0072-8

Langmead B, Salzberg SL. 2012. Fast gapped-read alignment with Bowtie 2. Nat Methods 9:357–359. doi:10.1038/nmeth.1923

Langmead B, Wilks C, Antonescu V, Charles R. 2019. Scaling read aligners to hundreds of threads on general-purpose processors. Bioinformatics 35:421–432. doi:10.1093/bioinformatics/bty648

Li H. 2018. Minimap2: Pairwise alignment for nucleotide sequences. Bioinformatics 34:3094–3100. doi:10.1093/bioinformatics/bty191

Li H, Handsaker B, Wysoker A, Fennell T, Ruan J, Homer N, Marth G, Abecasis G, Durbin R, 1000 Genome Project Data Processing Subgroup. 2009. The sequence Alignment/Map format and SAMtools. Bioinformatics 25:2078–2079. doi:10.1093/bioinformatics/btp352

Lohe AR, Roberts PA. 2000. Evolution of DNA in heterochromatin: The *Drosophila melanogaster* sibling species subgroup as a resource. Genetica 109:125–130. doi:10.1023/A:1026588217432

Long EO, Dawid IB. 1980. Repeated genes in eukaryotes. Annu Rev Biochem 49:727–764. doi:49.070180.003455

Long EO, Dawid IB. 1979. Expression of ribosomal DNA insertions in *Drosophila melanogaster*. Cell 18:1185–1196. doi:10.1016/0092-8674(79)90231-9

Lohse K, Mackintosh A, Vila R, Darwin Tree of Life Barcoding collective, Wellcome Sanger Institute Tree of Life programme, Wellcome Sanger Institute Scientific Operations: DNA Pipelines collective, Tree of Life Core Informatics collective, Darwin Tree of Life Consortium. 2021 The genome sequence of the European peacock butterfly, *Aglais io* (Linnaeus, 1758) [version 1; peer review: awaiting peer review]. Wellcome Open Res 6:258 doi:10.12688/wellcomeopenres.17204.1

Lu Y, Kochert GD, Isenhour DJ, Adang MJ. 1994. Molecular characterization of a strain-specific repeated DNA sequence in the fall armyworm *Spodoptera frugiperda* (Lepidoptera: Noctuidae). Insect Molecular Biology 3:123–130. doi:10.1111/j.1365-2583.1994.tb00159.x

Lukhtanov V. 2015. The blue butterfly *Polyommatus (Plebicula) atlanticus* (Lepidoptera, Lycaenidae) holds the record of the highest number of chromosomes in the non-polyploid eukaryotic organisms. Comparat Cytogenet 9:683–690. doi:10.3897/CompCytogen.v9i4.5760

Mahendran B, Acharya C, Dash R, Ghosh SK, Kundu SC. 2006. Repetitive DNA in tropical tasar silkworm *Antheraea mylitta*. Gene 370:51–57. doi:10.1016/j.gene.2005.11.010

Mandrioli M, Manicardi GC, Marec F. 2003. Cytogenetic and molecular characterization of the MBSAT1 satellite DNA in holokinetic chromosomes of the cabbage moth, *Mamestra brassicae* (Lepidoptera). Chromosome Res 11:51–56. doi:10.1023/a:1022058032217

Martí E, Milani D, Bardella VB, Albuquerque L, Song H, Palacios-Gimenez OM, Cabral-de-Mello DC. 2021. Cytogenomic analysis unveils mixed molecular evolution and recurrent chromosomal rearrangements shaping the multigene families on *Schistocerca* grasshopper genomes. Evolution 75:2027–2041. doi:10.1111/evo.14287

McKinlay A, Fultz D, Wang F, Pikaard CS. 2021. Targeted enrichment of rRNA gene tandem arrays for ultra-long sequencing by selective restriction endonuclease digestion. Front Plant Sci 12:656049. doi:10.3389/fpls.2021.656049

Mediouni J, Fuková I, Frydrychová R, Dhouibi MH, Marec F. 2004. Karyotype, sex chromatin and sex chromosome differentiation in the carob moth, *Ectomyelois ceratoniae* (Lepidoptera: Pyralidae). Caryologia 57:184–194. doi:10.1080/00087114.2004.10589391

Muirhead CA, Presgraves DC. 2021. Satellite DNA-mediated diversification of a sex-ratio meiotic drive gene family in *Drosophila*. Nat Ecol Evol 5:1604–1612. doi:10.1038/s41559-021-01543-8

Nelson JO, Slicko A, Yamashita YM. 2021. The retrotransposon R2 maintains *Drosophila* ribosomal DNA repeats. doi:10.1101/2021.07.12.451825

Nguyen P, Sahara K, Yoshido A, Marec F. 2010. Evolutionary dynamics of rDNA clusters on chromosomes of moths and butterflies (Lepidoptera). Genetica 138:343–354. doi:10.1007/s10709-009-9424-5

Noller HF, Lancaster L, Mohan S, Zhou J. 2017. Ribosome structural dynamics in translocation: yet another functional role for ribosomal RNA. Quart Rev Biophys 50:e12. doi:10.1017/S0033583517000117

Novák P, Neumann P, Macas J. 2010. Graph-based clustering and characterization of repetitive sequences in next-generation sequencing data. BMC Bioinformatics 11:378. doi:10.1186/1471-2105-11-378

Novák P, Neumann P, Pech J, Steinhaisl J, Macas J. 2013. RepeatExplorer: a Galaxy-based web server for genome-wide characterization of eukaryotic repetitive elements from next-generation sequence reads. Bioinformatics 29:792–793. doi:10.1093/bioinformatics/btt054

Palacios-Gimenez OM, Castillo ER, Martí DA, Cabral-de-Mello DC. 2013. Tracking the evolution of sex chromosome systems in Melanoplinae grasshoppers through chromosomal mapping of repetitive DNA sequences. BMC Evol Biol 13:167. doi:10.1186/1471-2148-13-167

Perumal S, Waminal NE, Lee Jonghoon, Lee Junki, Choi B-S, Kim HH, Grandbastien M-A, Yang T-J. 2017. Elucidating the major hidden genomic components of the A, C, and AC genomes and their influence on *Brassica* evolution. Sci Rep 7:17986. doi:10.1038/s41598-017-18048-9

Poletto AB, Ferreira IA, Cabral-de-Mello DC, Nakajima RT, Mazzuchelli J, Ribeiro HB, Venere PC, Nirchio M, Kocher TD, Martins C. 2010. Chromosome differentiation patterns during cichlid fish evolution. BMC Genet 11:50. doi:10.1186/1471-2156-11-50

Prins JD, Saitoh K. 2003. Karyology and sex determination In: Kükenthal W, editor. Band 4: Arthropoda, 2 Hälfte: Insecta, Lepidoptera, Moths and Butterflies, Teilband/Part 36, Vol 2: Morphology, Physiology, and Development. Berlin, Boston: DE GRUYTER. doi:10.1515/9783110893724.449

Prokopowich CD, Gregory TR, Crease TJ. 2003. The correlation between rDNA copy number and genome size in eukaryotes. Genome 46:48–50. doi:10.1139/g02-103

Proux-Wéra E, Byrne KP, Wolfe KH. 2013. Evolutionary mobility of the ribosomal DNA array in yeasts. Genome Biol Evol 5:525–531. doi:10.1093/gbe/evt022

Provazníková I, Hejníčková M, Visser S, Dalíková M, Carabajal Paladino LZ, Zrzavá M, Voleníková A, Marec F, Nguyen P. 2021. Large-scale comparative analysis of cytogenetic markers across Lepidoptera. Sci Rep 11:12214. doi:10.1038/s41598-021-91665-7

Raskina O, Barber JC, Nevo E, Belyayev A. 2008. Repetitive DNA and chromosomal rearrangements: Speciation-related events in plant genomes. Cytogenet Genome Res 120:351–357. doi:10.1159/000121084

Raskina O, Belyayev A, Nevo E. 2004. Activity of the *En/Spm*-like transposons in meiosis as a base for chromosome repatterning in a small, isolated, peripheral population of *Aegilops speltoides* Tausch. Chromosome Res 12:153–161. doi:10.1023/B:CHRO.0000013168.61359.43

Robinson R. 1971. Lepidoptera Genetics. Oxford: Pergamon Press.

Roiha H, Miller J, Woods L. 1981. Arrangements and rearrangements of sequences flanking the two types of rDNA insertion in *D. melanogaster*. Nature 290:749–754. doi:10.1038/290749a0

Ruiz-Herrera A, Nergadze SG, Santagostino M, Giulotto E. 2008. Telomeric repeats far from the ends: Mechanisms of origin and role in evolution. Cytogenet Genome Res 122:219–228. doi:10.1159/000167807

Scacchetti PC, Alves de Oliveira JC, Utsunomia R, Claro FL, de Almeida Toledo LF, Oliveira C, Foresti F. 2012. Molecular characterization and physical mapping of two classes of 5S rDNA in the genomes of *Gymnotus sylvius* and *G. inaequilabiatus* (Gymnotiformes, Gymnotidae). Cytogenet Genome Res 136:131–137. doi:10.1159/000335658

Scali V, Coluccia E, Deidda F, Lobina C, Deiana AM, Salvadori S. 2016. Co-localization of ribosomal and telomeric sequences in *Leptynia* (Insecta: Phasmatodea). Ital J Zool 83:285–290. doi:10.1080/11250003.2016.1219403

Sember A, Bohlen J, Šlechtová V, Altmanová M, Pelikánová Š, Ráb P. 2018. Dynamics of tandemly repeated DNA sequences during evolution of diploid and tetraploid botiid loaches (Teleostei: Cobitoidea: Botiidae). PLoS ONE 13:e0195054. doi:10.1371/journal.pone.0195054

Šíchová J, Nguyen P, Dalíková M, Marec F. 2013. Chromosomal evolution in tortricid moths: Conserved karyotypes with diverged features. PLoS ONE 8:e64520. doi:10.1371/journal.pone.0064520

Šíchová J, Ohno M, Dincă V, Watanabe M, Sahara K, Marec F. 2016. Fissions, fusions, and translocations shaped the karyotype and multiple sex chromosome constitution of the northeast-Asian wood white butterfly, *Leptidea amurensis*. Biol J Linn Soc 118:457–471. doi:10.1111/bij.12756

Šíchová J, Voleníková A, Dincă V, Nguyen P, Vila R, Sahara K, Marec F. 2015. Dynamic karyotype evolution and unique sex determination systems in Leptidea wood white butterflies. BMC Evol Biol 15:89. doi:10.1186/s12862-015-0375-4

Silva DMZ de A, Pansonato-Alves JC, Utsunomia R, Araya-Jaime C, Ruiz-Ruano FJ, Daniel SN, Hashimoto DT, Oliveira C, Camacho JPM, Porto-Foresti F, Foresti F. 2014. Delimiting the origin of a B chromosome by FISH mapping, chromosome painting and DNA sequence analysis in *Astyanax paranae* (Teleostei, Characiformes). PLoS ONE 9:e94896. doi:10.1371/journal.pone.0094896

Sims J, Sestini G, Elgert C, von Haeseler A, Schlögelhofer P. 2021. Sequencing of the *Arabidopsis* NOR2 reveals its distinct organization and tissue-specific rRNA ribosomal variants. Nat Commun 12:387. doi:10.1038/s41467-020-20728-6

Smit AFA, Hubley R, Green P. 2013. RepeatMasker Open-4.0.

Sochorová J, Garcia S, Gálvez F, Symonová R, Kovařík A. 2018. Evolutionary trends in animal ribosomal DNA loci: introduction to a new online database. Chromosoma 127:141–150. doi:10.1007/s00412-017-0651-8

Sproul JS, Barton LM, Maddison DR. 2020. Repetitive DNA profiles reveal evidence of rapid genome evolution and reflect species boundaries in ground beetles. Systematic Biology 69:1137–1148. doi:10.1093/sysbio/syaa030

Symonová R. 2019. Integrative rDNAomics—Importance of the oldest repetitive fraction of the eukaryote genome. Genes 10:345. doi:10.3390/genes10050345

Symonová R, Majtánová Z, Sember A, Staaks GB, Bohlen J, Freyhof J, Rábová M, Ráb P. 2013. Genome differentiation in a species pair of coregonine fishes: an extremely rapid speciation driven by stress-activated retrotransposons mediating extensive ribosomal DNA multiplications. BMC Evol Biol 13:42. doi:10.1186/1471-2148-13-42

Van Nieukerken EJ, Kaila L, Kitching IJ, Kristensen NP, Lees DC, Minet J, Mitter C, Mutanen M, Regier JC, Simonsen TJ, Wahlberg N, Yen S-H, Zahiri R, Adamski D, Baixeras J, Bartsch D, Bengtsson BÅ, Brown JW, Bucheli SR, Davis DR, Prins JD, Prins WD, Epstein ME, Gentili-Poole P, Gielis C, Hättenschwiler P, Hausmann A, Holloway JD, Kallies A, Karsholt O, Kawahara AY, Koster SJC, Kozlov MV, Lafontaine JD, Lamas G, Landry J-F, Lee S, Nuss M, Park K-T, Penz C, Rota J, Schintlmeister A, Schmidt BC, Sohn J-C, Solis MA, Tarmann GM, Warren AD, Weller S, Yakovlev RV, Zolotuhin VV, Zwick A. 2011. Order Lepidoptera Linnaeus, 1758. In: Zhang, Z.-Q. (Ed.) Animal biodiversity: An outline of higher-level classification and survey of taxonomic richness. Zootaxa 3148:212. doi:10.11646/zootaxa.3148.1.41

Van’t Hof AE, Nguyen P, Dalíková M, Edmonds N, Marec F, Saccheri IJ. 2013. Linkage map of the peppered moth, *Biston betularia* (Lepidoptera, Geometridae): a model of industrial melanism. Heredity 110:283–295. doi:10.1038/hdy.2012.84

Věchtová P, Dalíková M, Sýkorová M, Žurovcová M, Füssy Z, Zrzavá M. 2016. CpSAT-1, a transcribed satellite sequence from the codling moth, *Cydia pomonella*. Genetica 144:385–395. doi:10.1007/s10709-016-9907-0

Vershinina AO, Anokhin BA, Lukhtanov VA. 2015. Ribosomal DNA clusters and telomeric (TTAGG)n repeats in blue butterflies (Lepidoptera, Lycaenidae) with low and high chromosome numbers. CCG 9:161–171. doi:10.3897/CompCytogen.v9i2.4715

Vítková M, Fuková I, Kubíčková S, Marec F. 2007. Molecular divergence of the W chromosomes in pyralid moths (Lepidoptera). Chromosome Res 15:917–930. doi:10.1007/s10577-007-1173-7

Vondrak T, Oliveira L, Novák P, Koblížková A, Neumann P, Macas J. 2021. Complex sequence organization of heterochromatin in the holocentric plant *Cuscuta europaea* elucidated by the computational analysis of nanopore reads. Comput Struct Biotechnol 19:2179–2189. doi:10.1016/j.csbj.2021.04.011

Wolf KW, Novak K, Marec F. 1997. Kinetic organization of metaphase I bivalents in spermatogenesis of Lepidoptera and Trichoptera species with small chromosome numbers. Heredity 79:35–143.

Wu S, Xiong J, Yu Y. 2015. Taxonomic resolutions based on 18S rRNA genes: A case study of subclass Copepoda. PLoS ONE 10:e0131498. doi:10.1371/journal.pone.0131498

Xiong Y, Burke WD, Jakubczak JL, Eickbush TH. 1988. Ribosomal DNA insertion elements R1Bm and R2Bm can transpose in a sequence specific manner to locations outside the 28S genes. Nucleic Acids Res 16:10561–10573. doi:10.1093/nar/16.22.10561

Yano CF, Merlo MA, Portela-Bens S, Cioffi M de B, Bertollo LAC, Santos-Júnior CD, Rebordinos L. 2020. Evolutionary dynamics of multigene families in *Triportheus* (Characiformes, Triportheidae): A transposon mediated mechanism? Front Mar Sci 7:6. doi:10.3389/fmars.2020.00006

Yasukochi Y, Tanaka-Okuyama M, Kamimura M, Nakano R, Naito Y, Ishikawa Y, Sahara K. 2011. Isolation of BAC clones containing conserved genes from libraries of three distantly related moths: A useful resource for comparative genomics of Lepidoptera. J Biomed Biotechnol 2011:1–6. doi:10.1155/2011/165894

Zhou J, Eickbush MT, Eickbush TH. 2013. A Population genetic model for the maintenance of R2 retrotransposons in rRNA gene loci. PLoS Genet 9:e1003179. doi:10.1371/journal.pgen.1003179

Zrzavá M, Hladová I, Dalíková M, Šíchová J, Õunap E, Kubíčková S, Marec F. 2018. Sex chromosomes of the iconic moth *Abraxas grossulariata* (Lepidoptera, Geometridae) and its congener *A. sylvata*. Genes 9:279. doi:10.3390/genes9060279

