## Supplemental figures for "The role of repetitive DNA in re-patterning of major rDNA clusters in Lepidoptera"

**Table S3:** Summary of coverage analysis of individual elements in rDNA units in *H. humuli*, *A. urticae*, and *I. io*

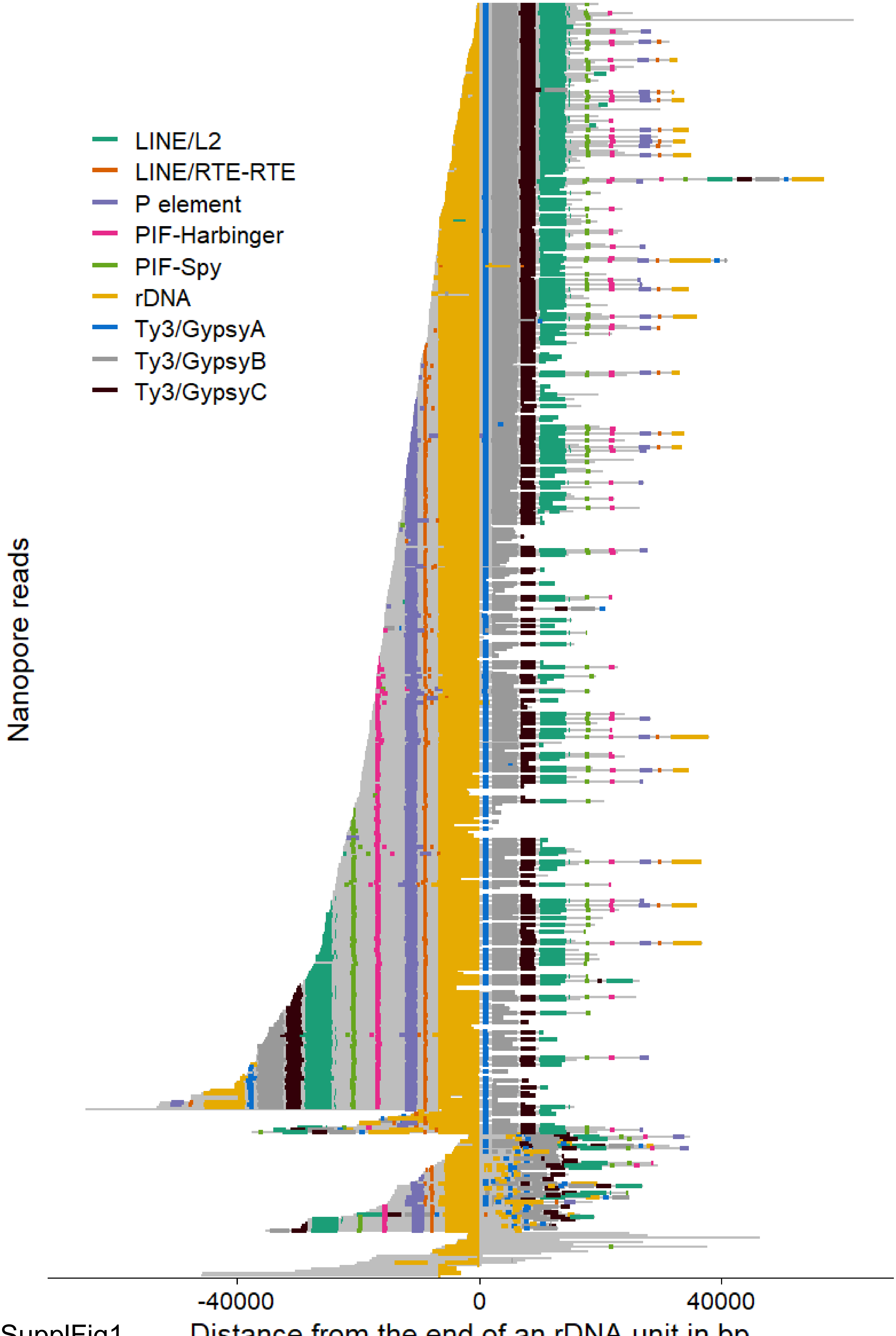
LINE/L2
LINE/RTE-RTE
P element
PIF-Harbinger
PIF-Spy
rDNA
Ty3/GypsyA
Ty3/GypsyB
Ty3/GypsyC
Nanopore reads
-40000
0
40000
Distance from the end of an rDNA unit in bp

SupplFig1

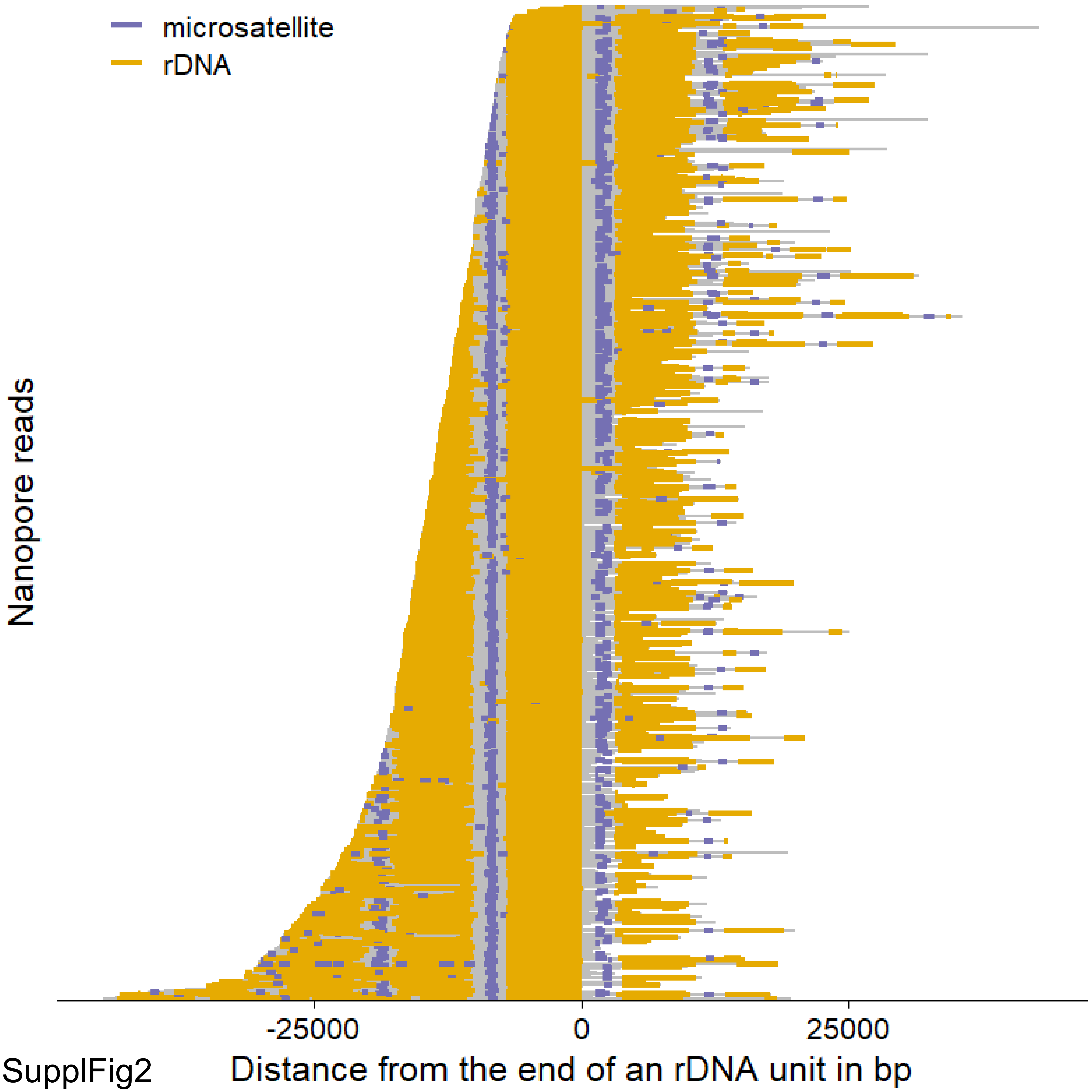

microsatellite
rDNA
Nanopore reads
-25000
0
25000
Distance from the end of an rDNA unit in bp

SupplFig2

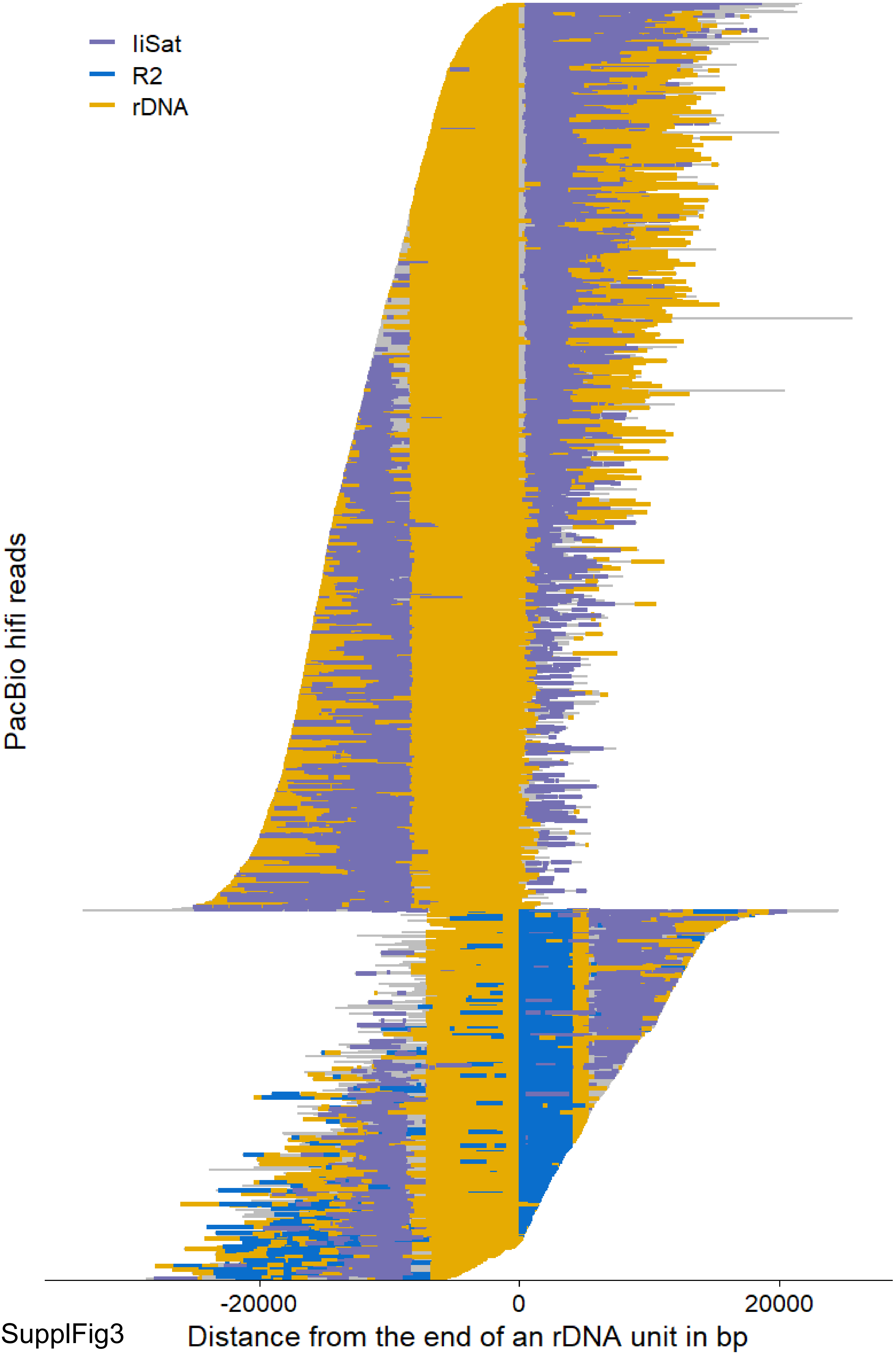
liSat
R2
rDNA
PacBio hifi reads
-20000
0
20000
Distance from the end of an rDNA unit in bp

SupplFig3

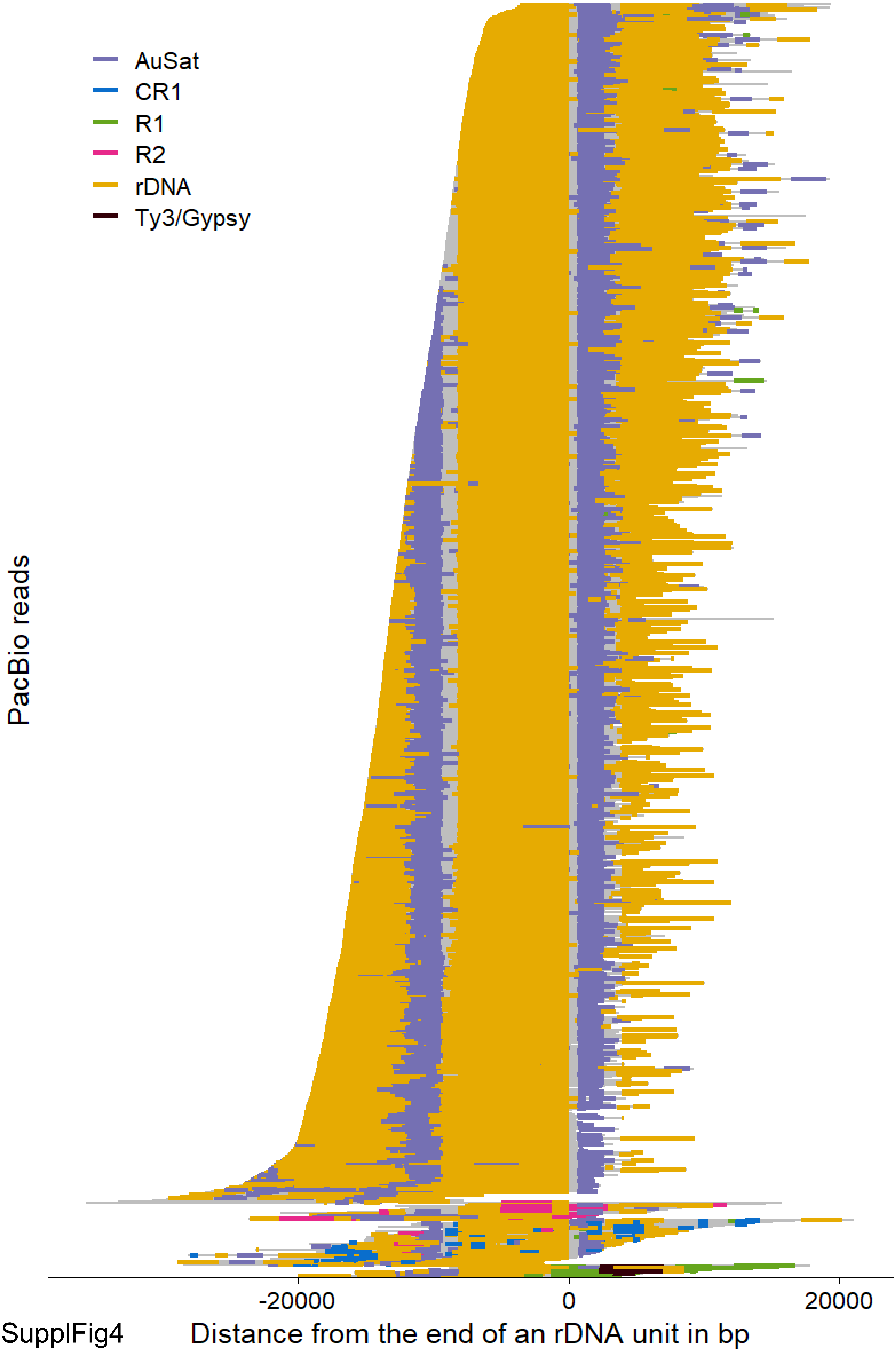

AuSat
CR1
R1
R2
rDNA
Ty3/Gypsy
PacBio reads
-20000
0
20000
Distance from the end of an rDNA unit in bp


SupplFig4

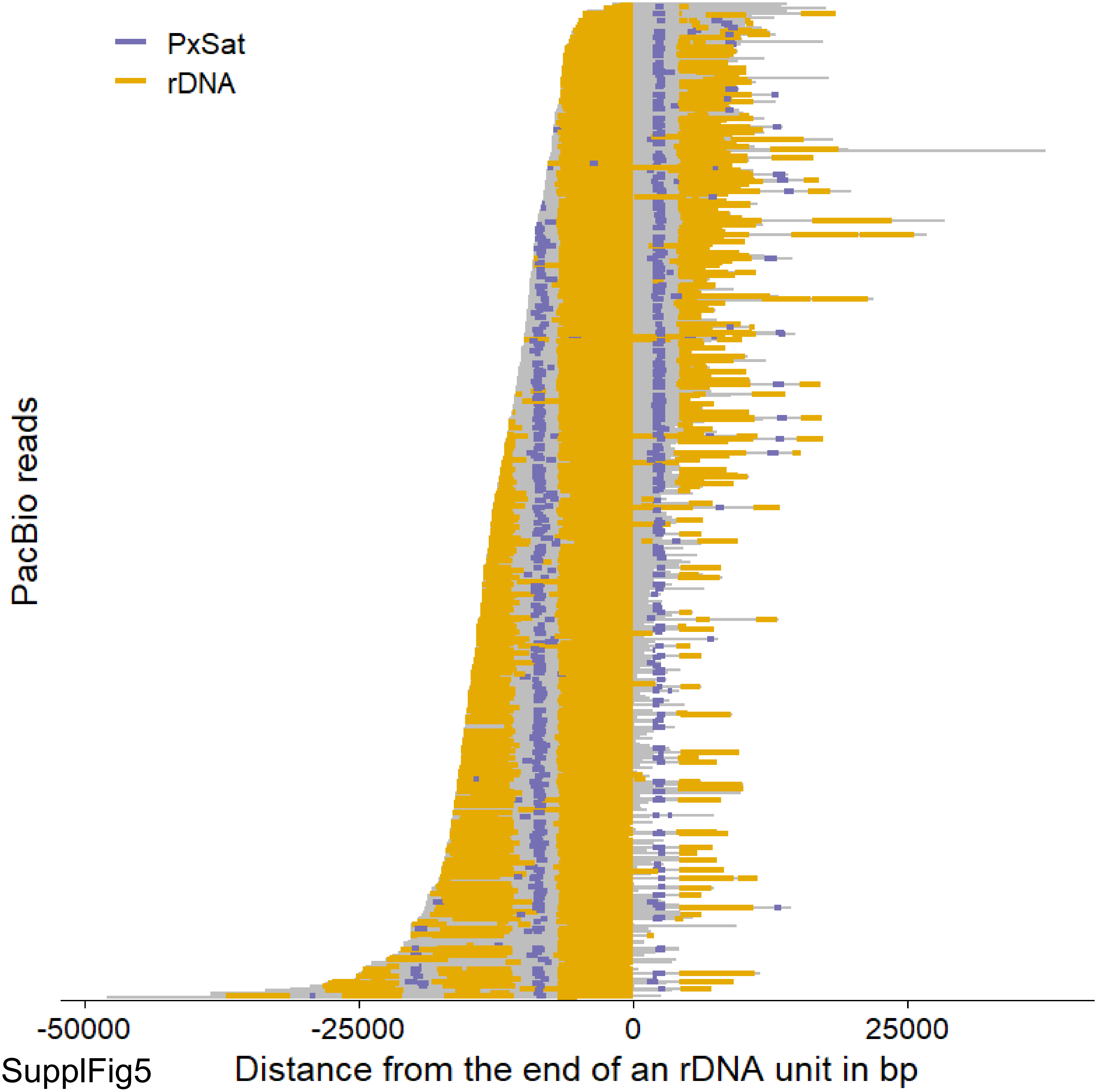

PxSat
rDNA
PacBio reads
-50000
-25000
0
25000
Distance from the end of an rDNA unit in bp


SupplFig5

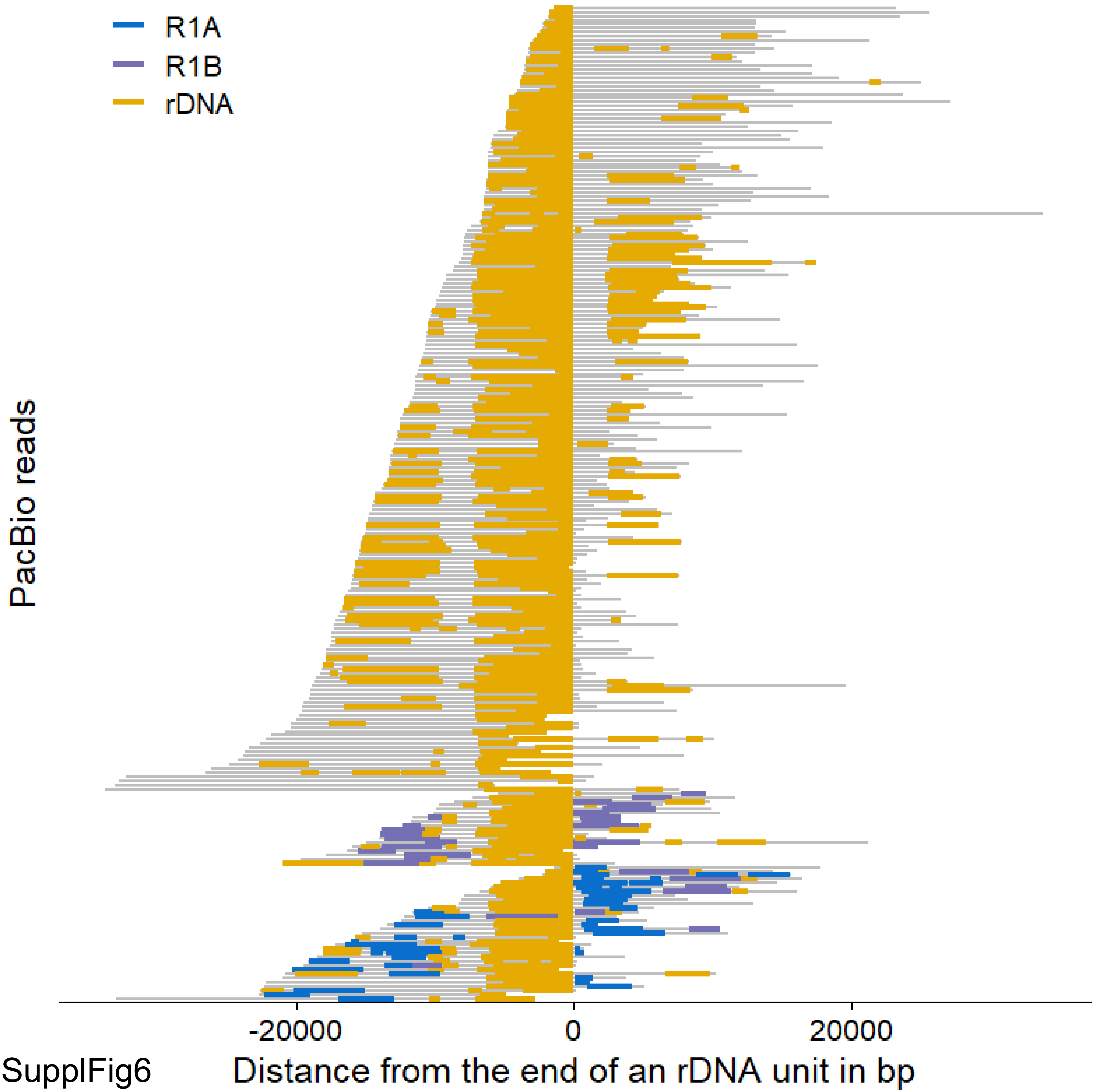
R1A
R1B
rDNA
PacBio reads
-20000
0
20000
Distance from the end of an rDNA unit in bp

SupplFig6

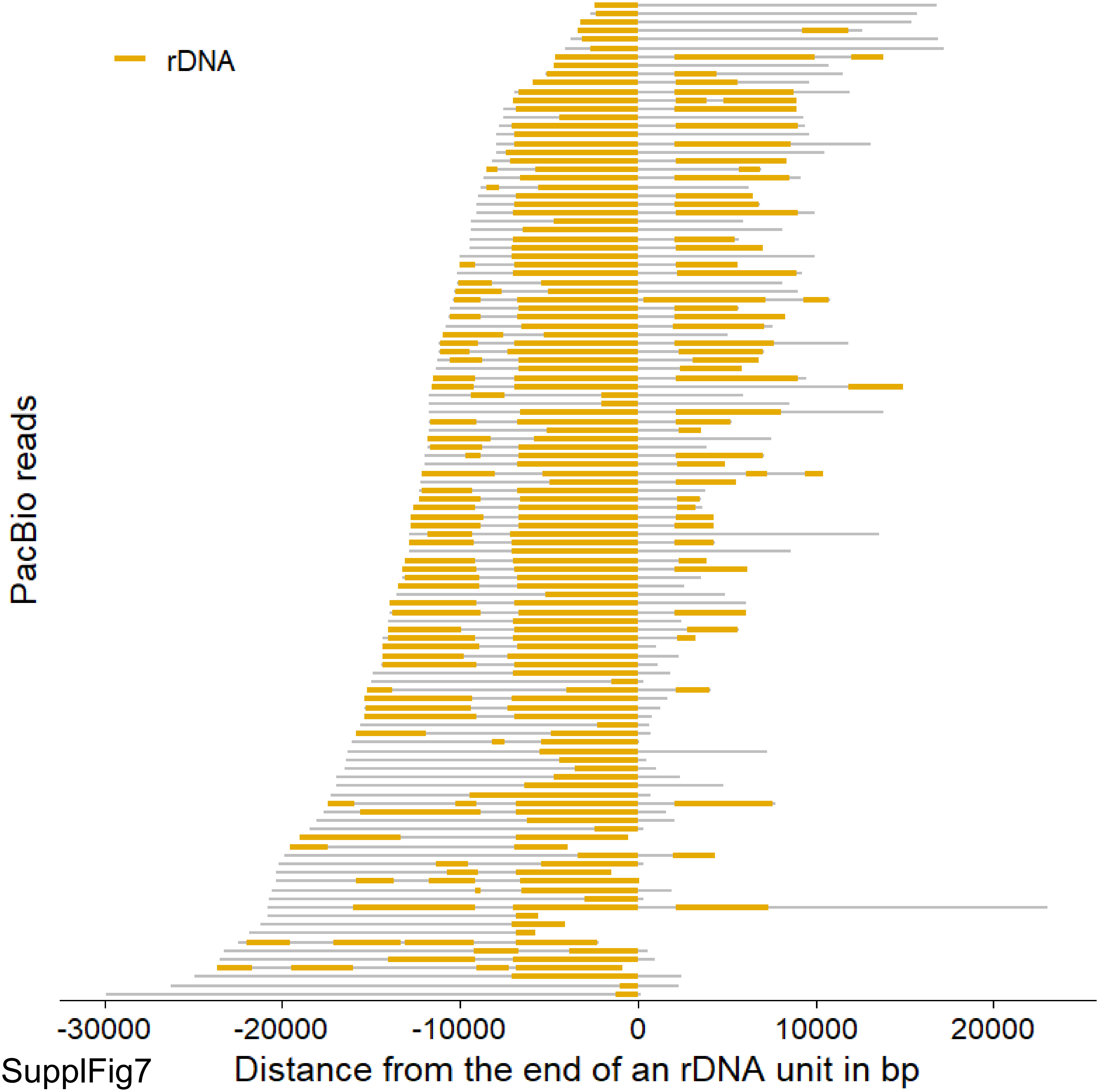
rDNA
PacBio reads
-30000
-20000
-10000
0
10000
20000
Distance from the end of an rDNA unit in bp

SupplFig7

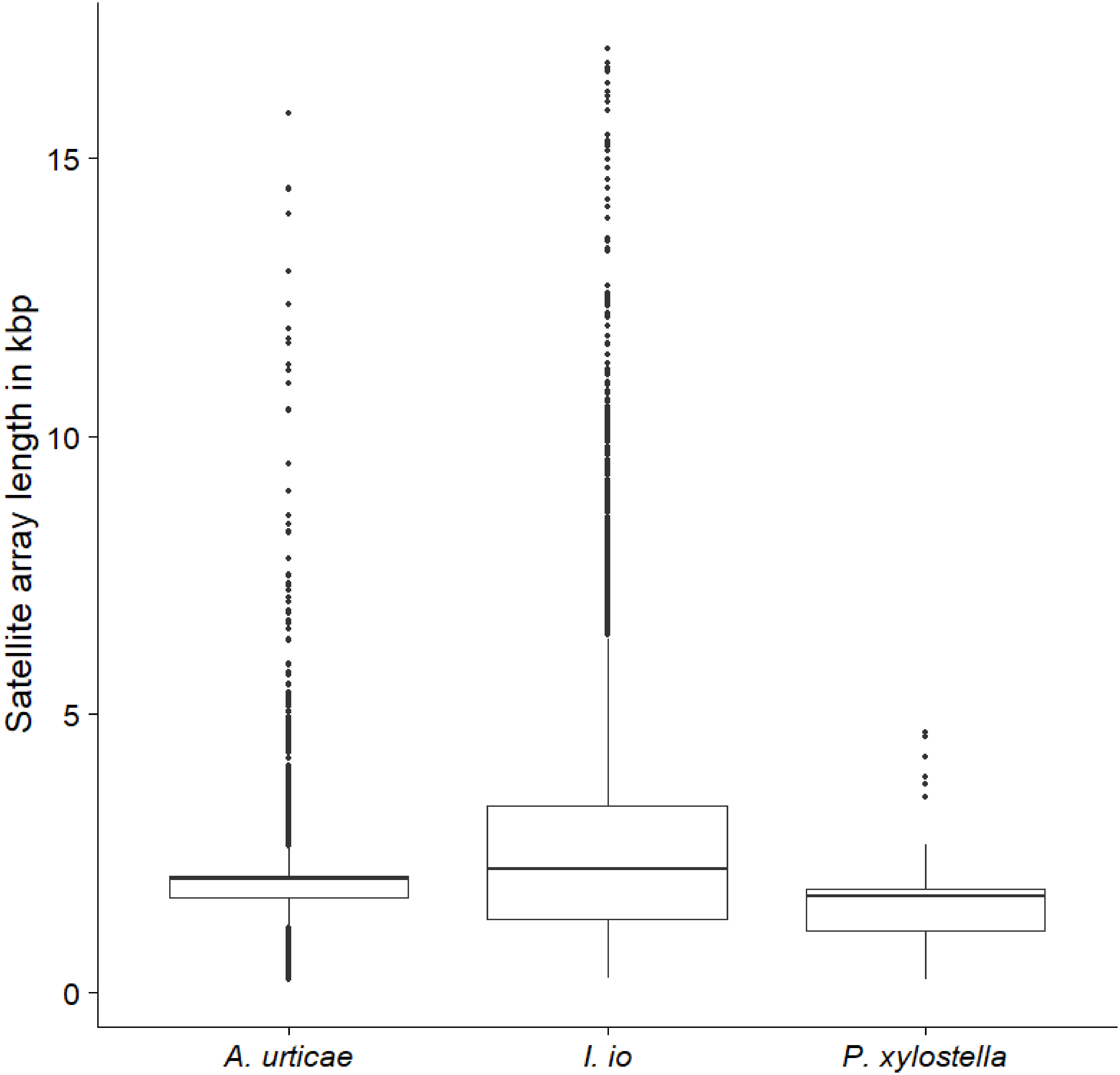

Satellite array length in kbp
15
10
5
0
A. urticae
I. io
P. xylostella


SupplFig8

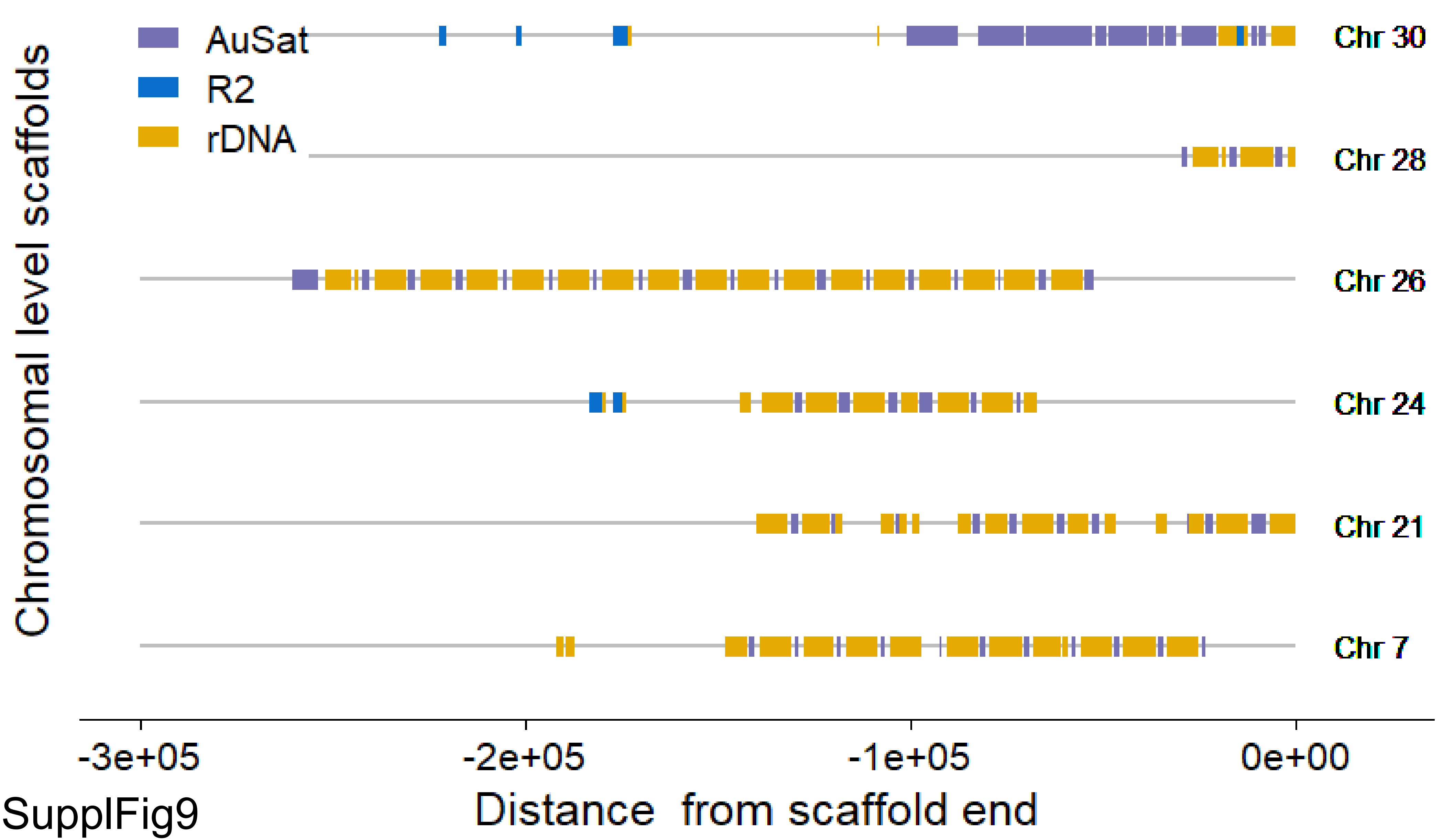
AuSat
R2
rDNA
Chr 30
Chr 28
Chr 26
Chr 24
Chr 21
Chr 7
Chromosomal level scaffolds
-3e+05
-2e+05
-1e+05
0e+00
Distance from scaffold end

SupplFig9
